# Context-dependent translation inhibition as a novel oncology therapeutic modality

**DOI:** 10.1101/2025.01.09.632223

**Authors:** Paige D. Diamond, Paul V. Sauer, Mikael Holm, Canessa J. Swanson-Swett, Lucas Ferguson, Natalie M. Bratset, Grant W. Wienker, Justin Seiwert Sim, Hailey K. Adams, Lillian Kenner, Margot Meyers, David Gygi, Zef A. Könst, Sogole Sami Bahmanyar, Lawrence G. Hamann, Anthony P. Schuller

**Affiliations:** Interdict Bio, 201 Haskins Way, South San Francisco, CA 94080; Interdict Bio, 201 Haskins Way, Suite 507, South San Francisco, CA 94080

**Keywords:** translation inhibitor, interdictor, ribosome, cryogenic-electron microscopy (cryo-EM), ribotoxic stress response, cancer, MYC

## Abstract

Inhibitors of protein synthesis, including anisomycin, homoharringtonine, and other natural products bind in the peptidyl-transferase center (PTC) of the eukaryotic ribosome to inhibit translation. Recent work has demonstrated that some PTC-binding antibiotics act in a sequence-selective manner, inhibiting translation elongation at specific amino acids while the polypeptide is engaged in the PTC. However, this phenomenon has yet to be documented for compounds that inhibit translation by the human ribosome. Here we use structure-based design to guide synthesis of molecules called interdictors that bind to the human ribosome PTC and act in a context-selective manner to inhibit translation elongation. Using ribosome profiling, in combination with *in vitro* biochemistry and cryo-electron microscopy, we characterize the context selectivity of unique analogues and observe their preferred interactions with nascent chain residues with complementary properties. Furthermore, we present a structure for an interdictor bound to a portion of the MYC protein at ∼ 1.9 Å resolution and identify resulting structural rearrangements in both the nascent chain and ribosomal RNA. In cells, we document how these compounds differentially impact the ribotoxic stress response pathway which monitors ribosome collisions and can trigger apoptosis. Finally, we confirm their tumor growth inhibition activity after oral dosing in cell line derived xenografts in mice using the MDA-MB-231 model for triple-negative breast cancer. Together, our data establish sequence-selective inhibition of translation as a novel small-molecule therapeutic modality for historically difficult to address cancers by targeting translation of oncogenic dependency factors in the human ribosome PTC.

## INTRODUCTION

Inhibitors of protein synthesis can impact any of the four key phases of mRNA translation by the ribosome: initiation, elongation, termination and recycling^1–3^. Translation elongation (as the ribosome incorporates successive amino acids into the nascent protein) offers unique features for selective inhibition deriving from the diversity of properties of amino acids that comprise the newly synthesized polypeptide. While several natural product inhibitors of eukaryotic translation elongation have been identified and characterized^2,4–7^, only one has progressed to clinical use. Homoharringtonine (HHT) is FDA approved as an injectable second line treatment for chronic myeloid leukemia (CML) and continues to be evaluated clinically for other hematologic oncology indications^8,9^. However, HHT lacks any potential for selective target translation inhibition as it indiscriminately inhibits formation of the first peptide bond across all mRNAs^10,11^.

Over the past decade, a combination of improved structural, biochemical, and genomic approaches has revealed that many natural products historically understood to be global inhibitors of elongation have context-or sequence-dependent activity^12–15^. In bacterial systems, these include the macrolide antibiotics that bind in the nascent polypeptide exit tunnel (NPET) and interact with the growing protein, such as erythromycin or telithromycin^12–15^, and antibiotics that bind in the peptidyl-transferase center (PTC) and interact with the most recently incorporated amino acids in the linear nascent polypeptide, such as chloramphenicol^14,16^ or the oxazolidinones (linezolid and radezolid^15,16^). There is only one documented drug-like molecule binding to the NPET, PF846, that has been reported to have any sequence preference^17^, and which inhibited the translation of proprotein convertase subtilisin/kexin type 9 (PCSK9) by binding adjacent to the human ribosome NPET; however, no inhibitors that bind to the PTC of the human ribosome have been documented to have sequence-selectivity. While the polypeptide can form secondary structure in the NPET^18^, the narrow steric constraints of the PTC permit only a strictly linear polypeptide conformation which would enable rational design of molecules that interact with select target peptides with high precision. The human ribosome PTC is therefore an attractive target for structure-based drug design to inhibit the synthesis of select protein targets, especially those that may lack accessible binding pockets or are intrinsically disordered and therefore considered “undruggable” by traditional methods^19^.

An exemplary target for which targeted inhibition of translation may constitute a powerful therapeutic approach is MYC, a master regulator of many gene expression programs that promote cancer cell growth and survival^20^. MYC is among the most commonly activated (∼70%) genes in human cancers^20,21^, has both very rapid protein and RNA turnover kinetics^22,23^, and is almost entirely unstructured^21^, rendering any occupancy-based approaches both extremely challenging and stoichiometrically untenable. Global translation inhibition by HHT or other natural products has been previously reported to kill cancer cells by reducing pro-oncogenic, short half-life proteins of high survival dependency such as MYC and MCL1 or to sensitize cells to other chemotherapeutic agents^24–26^. An alternate mechanism by which translation inhibition has been shown to impact cancer cell viability is mediated through stress response pathways that lead to apoptosis, as has been documented for anisomycin^27,28^, further confirming translation inhibition as a therapeutic strategy even in the absence of biochemical selectivity.

We therefore set out to develop inhibitors of human translation that bind to the ribosome PTC and slow elongation in a sequence-selective manner. Leveraging the pharmacophore of the natural product anisomycin as a template, we designed and synthesized novel small molecules that confer amino-acid sequence selectivity through unique moieties that project toward and engage with the linear polypeptide chain in the PTC. Using ribosome profiling, we characterized the individual stall-sequence preferences of each of these molecules. As designed, we observed that each analogue preferentially interacts with nascent chain peptide residues that have complementary physiochemical properties to moieties of the small molecule, highlighting target sequence tunability of this novel modality. To cross-validate our ribosome profiling data, we performed *in vitro* translation assays on luciferase reporters containing the identified stall motifs and observe a >15-fold elongation inhibition selectivity of these molecules for their respective preferred motifs. Using cryogenic electron microscopy (cryo-EM) of purified human ribosome nascent chain complexes, we then observed sequence-specific contacts between each of these inhibitors and the nascent polypeptide chain. Structural determination of a MYC-programmed ribosome nascent chain complex to 1.9 Å further elucidates specific interactions between the peptide and an interdictor and how the nascent polypeptide is structurally rearranged upon compound binding. Additionally, we verify the activity of these molecules in a panel of MYC-dependent cancer cell lines where both compounds inhibit cancer cell proliferation but differentially impact the ribotoxic stress pathway which is activated by ribosome collisions to control cell homeostasis^29^. Finally, we confirm the *in vivo* efficacy of an interdictor in the MDA-MB-231 human TNBC xenograft model in mice where we observe 80% tumor growth inhibition after oral dosing. Together our data lay the foundation for sequence-selective translation modulation by small molecule drugs as an emerging powerful modality to address historically undruggable targets with high validation.

## RESULTS

### Interdictors lead to unique context-dependent translation inhibition in human cells

We leveraged composite pharmacophore descriptors from anisomycin and other known PTC binders to computationally design novel small molecule sequence-selective translation inhibitors. Our design principles were rooted in maintaining the stabilizing interactions of the anisomycin (ANS) shared core with the rRNA^30^, while modularly grafting on unique substituents that extend toward the nascent polypeptide chain and form stabilizing interactions with the emerging amino acids (Figure 1A). IDB-001 includes a basic (*S*)-2-(pyrrolidin-2-yl)ethan-1-amine (distal pyrrolidine) substituent which we predicted to preferentially bind and form salt-bridges with aspartic acid and glutamic acid residues in the nascent polypeptide. In contrast, we hypothesized that IDB-002 with a distal non-polar 3-fluorobenzyl substituent would preferentially interact with hydrophobic side chains.

**Figure 1.**
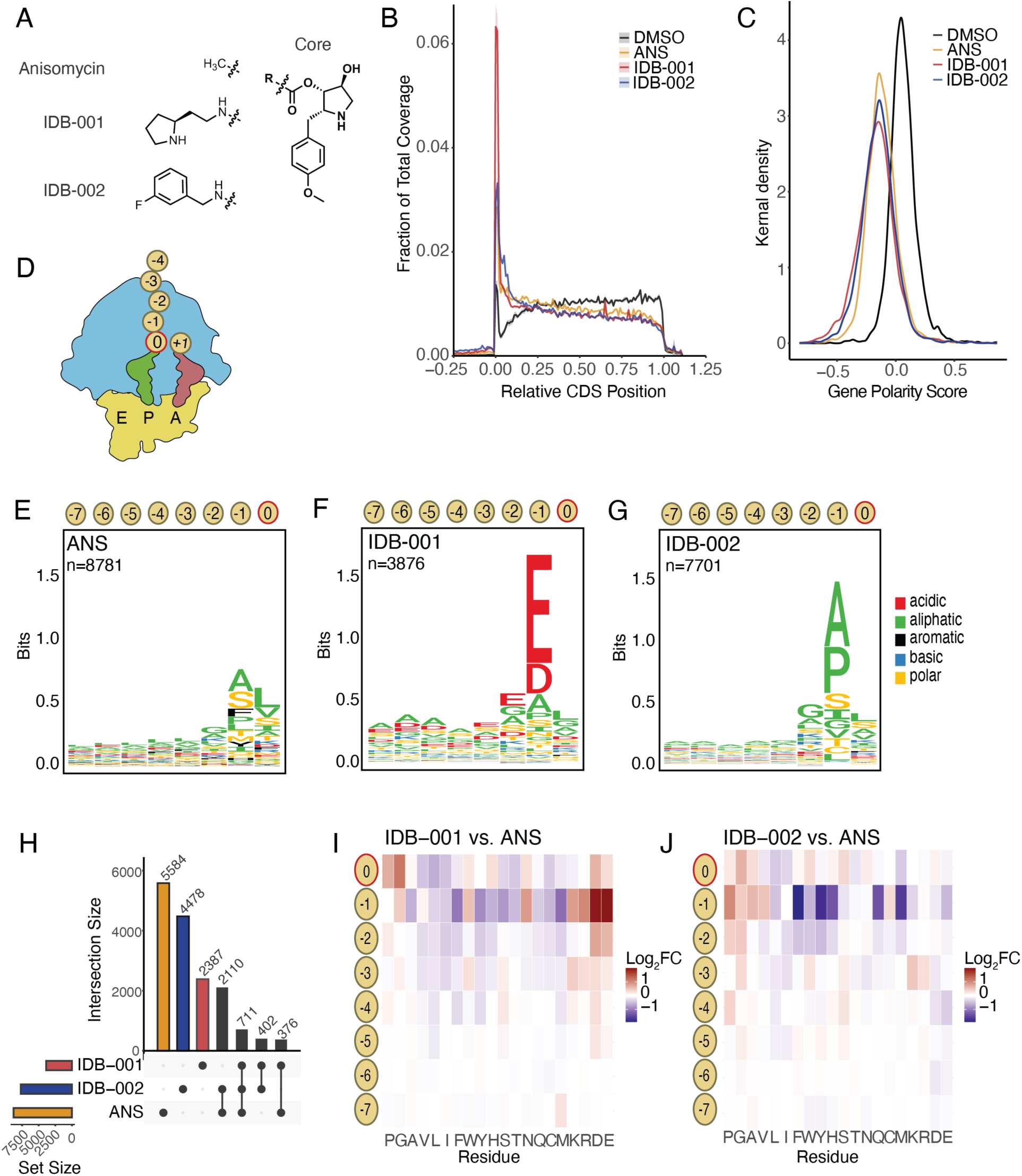
Interdictors lead to unique context-dependent translation inhibition in human cells. A. Structures of anisomycin (ANS) and interdictors IDB-001 and IDB-002. B. Metagene analysis of open-reading frames for IDB-001, IDB-002 (10 μM) and DMSO-treated samples. C. Kernel density plot of polarity scores across transcripts with sufficient coverage (> 0.1 read/codon). D. Schematic illustrating nascent polypeptide numbering convention where the P site is set to 0. E-G. Sequence logos for the nascent chain sequence from positions –7 to 0 for significantly enriched pause sites codons (with odds ratio > 1 and p-values < 0.05) for ANS-treated samples (E), IDB-001 treatment (F), or IDB-002 (G). N corresponds to the number of significantly enriched codon positions. H. UpSet plot of significantly enriched codon positions with differential ribosome occupancy across IDB-001, IDB-002 and ANS treatments. I-J. Position weighted matrix demonstrating changes in ribosome occupancy for each position in the nascent chain and amino acid residue identity between IDB-001 and ANS (I), or IDB-002 and ANS (J). Red color indicates a log_2_ fold change of +1, and blue –1.

To investigate the extent of context-dependent inhibition by these molecules in human cells, we performed ribosome profiling^11,31,32^ in the MYC-dependent triple-negative breast cancer (TNBC) cell line HCC-1143 treated with either ANS, IDB-001 or IDB-002. We first examined how these molecules globally affected ribosome occupancy across all coding sequences (CDS) in the transcriptome. We found a pronounced accumulation of footprints at the 5’ ends of the open reading frame (ORF), coupled with a decreased footprint density near the 3’ ends compared to DMSO-treated controls (Figure 1B, Extended Data Figure 1A-B), confirming compound treatment resulted in a defect in elongation^31,33–35^. Additionally, we observed different levels of ribosome coverage at the start codon itself between the two compounds, indicating that these molecules differ in their ability to inhibit the first round of elongation. We next sought to determine whether these global changes in ribosome coverage were uniform across all transcripts or driven by specific mRNAs. To this end, we developed a polarity score for each transcript that reflected the ribosome coverage at each codon, weighted by the position of that codon within the CDS^34^. Polarity scores range from –1, indicating that all ribosomes were positioned at the start codon, to +1, indicating that they were entirely clustered at the stop codon. In drug-treated cells, for both interdictors and ANS, polarity scores generally shifted towards negative values, indicating that the majority of transcripts (>95%) exhibited a redistribution of ribosomes towards the 5’ end (Figure 1C), again consistent with a strong defect in elongation^34^.

As interdictors are designed to stall translation in a sequence-selective manner in the PTC, we shifted our focus to understand how nascent polypeptide sequence context in the PTC influenced changes in ribosome occupancy in response to compound treatment. To this end, we first identified codons that had differential ribosome occupancy relative to total transcript coverage in drugged versus DMSO-treated cells by a two-tailed Fisher’s exact test^36^. The nascent polypeptide sequences of significantly enriched pause sites were used to generate sequence logos^37^ for positions –7 to 0 in the nascent chain (the portion of the nascent polypeptide still engaged in the PTC), where 0 corresponds to the amino acid residue encoded by the codon occupying the P site (Figure 1D). Notably, in cells treated with ANS, we observed increased ribosome occupancy for footprints containing alanine and serine residues in the –1 polypeptide position (E-site Ala/Ser) (Figure 1E). In contrast, IDB-001 pause motifs were enriched for acidic residues (Asp/Glu) in the –1 position in agreement with its structure-based design principles (Figure 1F). IDB-002 displayed a similar profile to ANS though with a few notable features, including an increased preference for prolines and small hydrophobic residues (Ala, Val, Ile) in the –1 position, and a depletion of large hydrophobic residues (Phe, Tyr) at the same position (Figure 1G). While we observe some overlap in the pause motifs between compounds, each molecule tested had a substantial number of uniquely targeted positions (Figure 1H & Extended Data Figure 1C).

To compare IDB-001 and IDB-002 more directly to ANS and minimize the impact of the shared pharmacophore, we computed a position weighted matrix (PWM) of the nascent peptides from position –7 to 0 for compounds relative to ANS (Figure 1I-J). Unlike the motif analyses, this comparison included all ribosome footprints among ORFs which had sufficient coverage (greater than 0.1 read/codon) across all samples rather than only the differential pause sites. In the case of IDB-001, this differential PWM analysis revealed that in addition to the strong acidic signature at position –1, this compound displayed acidic preferences further up the nascent chain (Figure 1I). Compared to ANS, IDB-001 also shows a minor enrichment of positively charged (Arg, Lys) residues at this position. ANS strongly disfavors these residues, and this apparent enrichment with IDB-001 may be due to its weaker overall inhibition (Extended Data Figure 1D-F). Notably, this enrichment disappears when IDB-001 is compared to DMSO (Extended Data Figure 1D-F). When comparing IDB-002 to ANS, we could more readily observe a depletion of ribosome footprints containing bulky side chains (such as Phe, Tyr, Trp) in the –1 position (Figure 1J). Overall, these findings highlight the sequence-selective stalling effects of different interdictors, revealing both shared and unique amino acid targeting preferences for each compound.

### Interdictors engage nascent polypeptide motifs to elicit sequence-selective ribosome stalling

To more deeply characterize the sequence selectivity of each compound, we titrated each molecule and analyzed pause motifs identified by ribosome profiling across a gradient of drug concentrations (Extended Data Figure 2). To globally analyze nascent polypeptide motifs enriched by each molecule, we counted the occurrence of every tetrapeptide (amino acids –3 to 0, referred to as “4-mers”) in our ribosome footprinting data and performed differential expression analysis^38^. As anticipated, we found that the number of significant 4-mers decreased with decreasing concentrations of both compounds, though the major trends we identified at the highest concentration remained prevalent (Figure 2A-B, Extended Data Figure 2C-J). By generating sequence logos for all 4-mers with significant enrichment at each concentration, we identified additional sequence features that were not seen in our initial global analysis (Figure 1F-G). At the lowest concentrations (80 nM), we observed tetrapeptide motifs that represent the strongest stall feature for each molecule as they remain significantly enriched. In our analyses of these motifs on individual endogenous genes (*e.g., ANLN* and *H2BC17*), we observe a significant build-up of ribosomes at these sites, and a depletion of footprints afterward (Extended Data Figure 3), suggestive of a strong elongation stall.

**Figure 2.**
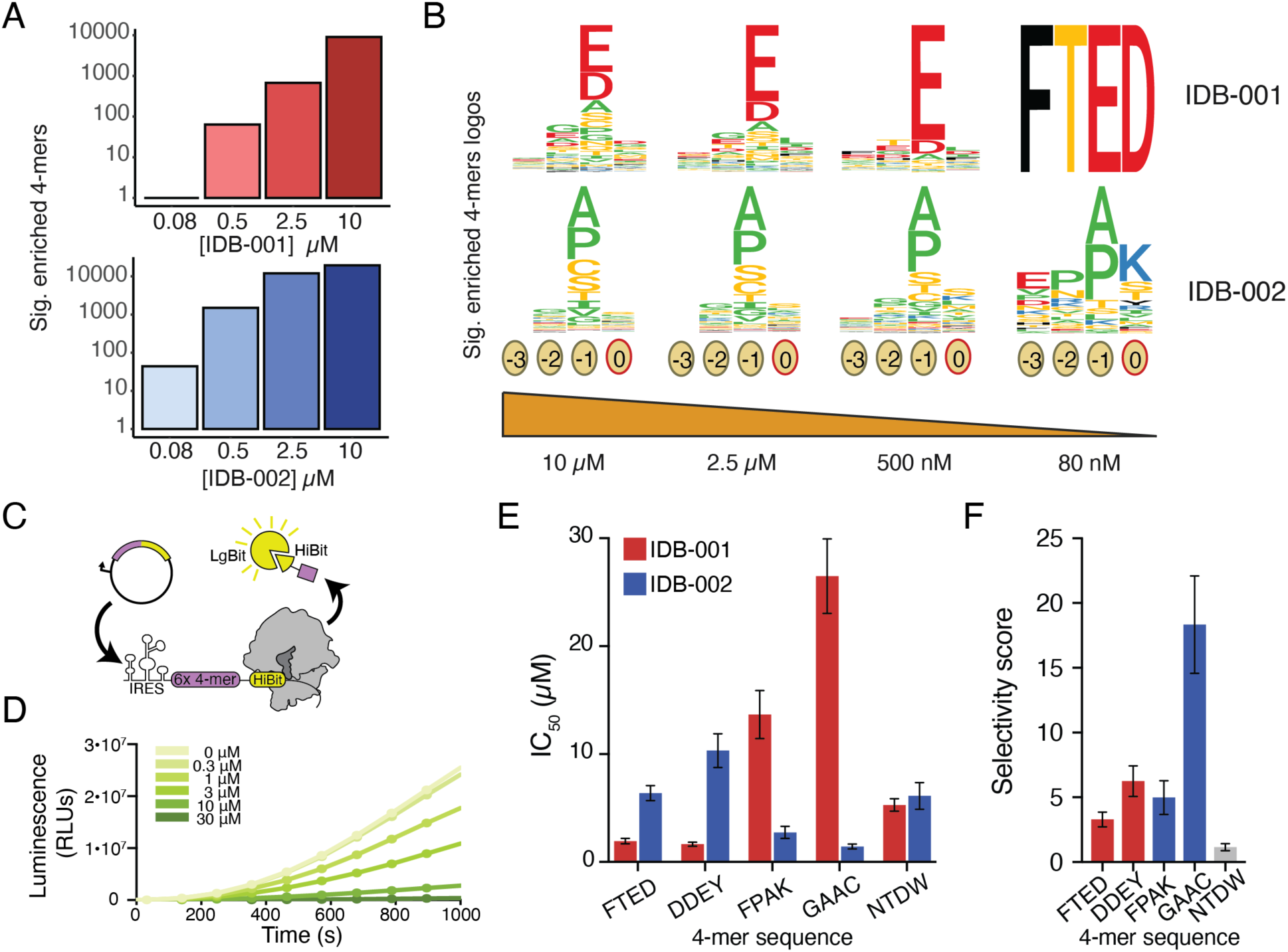
Interdictors engage select polypeptide motifs to inhibit translation kinetics. A. Bar graph illustrating significant number of enriched 4-mers identified by ribosome profiling titration of IDB-001 and IDB-002. B. Corresponding sequence logos for enriched 4-mers in (A). C. Schematic of the *in vitro* transcription-translation (IVTT) assay. After translation termination, the short reporter protein is released into solution where the HiBit-tag at the C-terminal end associates with LgBit protein to generate bioluminescence. The overall rate of protein synthesis is then quantified. D. Representative luminescence time traces for synthesis of the FTED reporter construct in the presence of increasing concentrations of IDB-001. E. IC_50_ values, determined from experiments such as in D, for each reporter treated with either IDB-001 (red) or IDB-002 (blue). F. Selectivity score (calculated as the ratio of IC_50_s for each reporter construct) where the color of the bar indicates the more potent compound, colored as in (E).

To further explore the selective elongation defect caused by our small molecules, we used an *in vitro* transcription-translation assay to probe the effect of these compounds on putative stalling or non-stalling peptide motifs. Using our ribosome occupancy data to predict the sensitivity of every 4-mer to IDB-001 and IDB-002, we generated a panel of reporter sequences for testing: two sequences predicted to be strongly inhibited by IDB-001 but not by IDB-002 (FTED & DDEY), two sequences predicted to be strongly inhibited by IDB-002 but not by IDB-001 (FPAK & GAAC), and one sequence that is not strongly preferred by either compound (NTDW). Each reporter contained six repeats of the identified pause motif 4-mer, followed by the 11 AA HiBit tag that produces luminescence when released from the ribosome (Figure 2C & Extended Data Figure 4A)^39,40^. Translation of these reporters, and inhibition by our compounds, could then be followed in real time using the IVTT system (Figure 2D).

We measured the effect of IDB-001 and IDB-002 on the steady-state rate of reporter protein synthesis and calculated IC_50_ values for each compound by analyzing translation rates versus compound concentration across an 11-point titration (Figure 2E & Extended Data Figure 4B). Comparing the IC_50_s of each compound on each reporter (Figure 2D), we observed 3-fold higher inhibitory activity of IDB-001 than IDB-002 on the FTED reporter, and 6-fold higher on the DDEY reporter (Figure 2F). Conversely, IDB-002 had a 5-fold stronger inhibitory effect on the FPAK reporter than IDB-001, and 18-fold stronger on the GAAC reporter (Figure 2F). As predicted, the NTDW reporter was equally inhibited by both compounds (Figure 2F). When comparing strongly inhibited compound-stall motif pairs and weakly inhibited compound-stall motif pairs for IDB-001, we find that there is at least a 15-fold range of biochemical specificity for nascent peptide sequences (Figure 2E & Extended Data Figure 4B, ratio of IC_50_ for DDEY versus GAAC). Together, these results connect the context-dependence we observe from ribosome profiling with a direct biochemical effect on translation rate, further confirming the sequence-selective mechanism of action of these compounds.

### Interdictors make sequence-specific interactions with the nascent chain

To better characterize the precise molecular interactions that underlie interdictor function and sequence preference, we used cryo-EM to determine structures of compound-bound human ribosome-nascent chain complexes (RNCs) stalled on enriched sequences identified by ribosome profiling. To obtain stalled complexes with the relevant 4-mers attached to the P-site tRNA, we programmed IVTT reactions with DNA encoding an mRNA reporter without a stop codon^41^ that ended in the 4-mer of interest (Extended Data Figure 5A). Purified RNCs carrying the 4-mers FTED or FPAK were then incubated with IDB-001 and IDB-002, respectively, before cryo-EM grid preparation.

The resulting structures showed that most tRNA-containing ribosomes were stalled in a classical, post-translocation state with a peptidyl-tRNA bound in the P site (Figure 3A). The map of the FTED-containing complex bound to IDB-001 reached a global resolution of 1.9 Å with a local resolution of 1.8 Å around the PTC, and the map of the FPAK-containing complex bound to IDB-002 reached a resolution of 2.2 Å with a local resolution of 2 Å in the PTC. In the PTC of both structures, there is clear density for each compound adjacent to the nascent chain (Figure 3A). Similar to ANS^30^, the core binds in a pocket formed by 28S rRNA bases including A3905, G3907, C4398, and the phosphate backbone of U4450, G4451, and U4452 (*E. coli* residues 2059, 2061, 2452, 2504, 2505, 2506 respectively) (Figure 3B). In addition, the core pyrrolidine is positioned to hydrogen bond with C4398 (Figure 3B). The 2-hydroxyl of the core pyrrolidine also hydrogen bonds with U4446 and coordinates a potassium ion (Figure 3C). Finally, the methoxyphenyl forms π-stacking interactions with C4398 as it extends towards the tRNA (Figure 3B), all consistent with previous reports^2,30^ and showing that the modification and addition of functional substituents to a shared pharmacophore of ANS does not interfere with ribosome binding.

**Figure 3.**
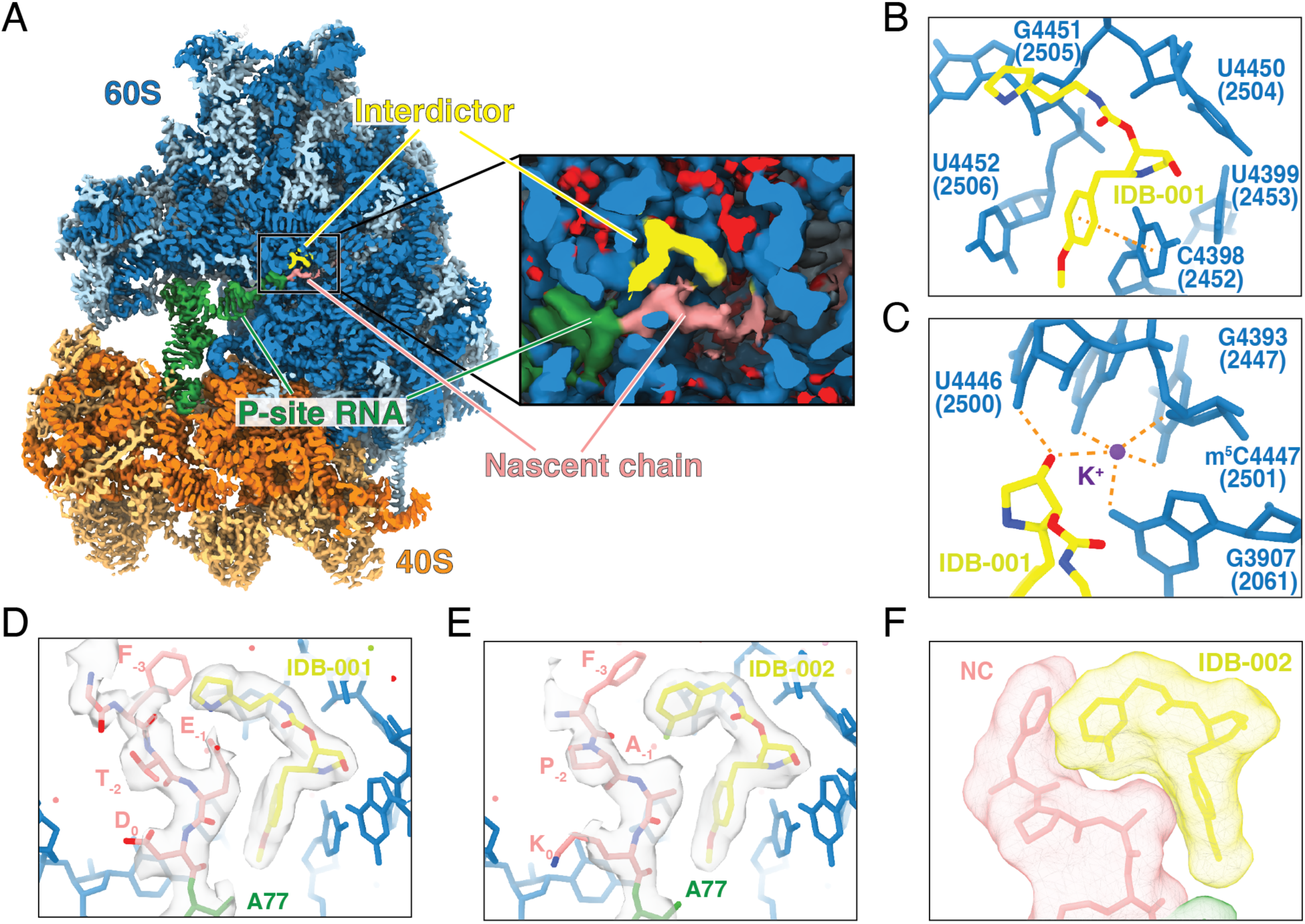
Structural analysis of interdictor-bound RNCs. A. Representative cryo-EM map of a stalled RNC with a tRNA in the P site, bound to IDB-001. Inset shows a closeup of the PTC containing the IDB-001 (yellow), P-site tRNA (green), nascent chain (salmon), rRNA (dark blue), and signal for ions and water molecules (red). B. The binding pocket of IDB-001 within the 28S rRNA of the ribosome. Residues are labeled with *H. sapiens* numbering and *E. coli* numbering in parentheses. π-stacking interaction between the benzyl group of IDB-001 and C4398 is indicated with orange dashed line. C. Hydrogen bonding network of IDB-001 to U4446 and a potassium ion. Bonds are indicated as orange dashed lines. D. Closeup of the FTED-containing RNC structure bound by IDB-001. E. Closeup of the FPAK-containing RNC structure bound by IDB-002. F. Surface representation of IDB-002 and the FPAK 4-mer to illustrate the shape complementarity.

In both maps, density for the nascent chain extends from the P-site tRNA past the small molecule and towards the exit tunnel (Figure 3D-E). Despite the inherently dynamic nature of the nascent chain in the exit tunnel, backbone and side chain positions of the four residues in contact with the compound (in the 0, –1, –2, and –3 positions) could be assigned. In each structure, the compound adopts a hook-like conformation in which the R-group substituent (Figure 1A) extends toward the nascent chain, resulting in engagement of a shape-complementary binding surface (Figure 3F). For IDB-001, the distal pyrrolidine binds in the pocket shaped by the peptide FTED where it forms a direct salt-bridge with E_-1_ and favorable electrostatic interactions with the backbone carbonyl of T_-2_ (Figure 3D, Extended Data Figure 6A-C). Additionally, the backbone methylene of E_-1_ is positioned to interact with the methoxybenzyl core of IDB-001 (Extended Data Figure 7A-B). Favorable cation-π interactions and van der Waals interactions can be made with F_-3_ (Extended Data Figure 6A-C), though the density for this residue is weak in our map suggesting it may be flexible. IDB-002, with its non-polar distal fluorophenyl moiety binds in the pocket formed by FPAK where the sidechains of A_-1_ and P_-2_ encapsulate the fluorophenyl substituent (Figure 3E, Extended Data Figure 6D-F). The A_-1_ side chain also is positioned to interact with the methoxybenzyl core of IDB-002 (Extended Data Figure 6D-E). In both cases, the interactions between the nascent polypeptide chain and compounds result from both steric and electrostatic complementarity of the distal substituents of IDB-001 and IDB-002. These structures show small molecule interdictors can engage in sequence-selective interactions, leveraging shape and charge complementarity, and validating our targeted structure-based design approach.

### Structural rearrangements upon binding of IDB-001 to a MYC peptide-containing RNC

To demonstrate the ability of these molecules to engage therapeutically relevant targets during their translation, we selected an acidic stretch within the oncogenic transcription factor MYC (D_251_SEEEQEDE_259_)^32^. Using the same biochemical approach as above, we generated ribosomes stalled on a MYC non-stop mRNA and incubated with and without IDB-001 before grid preparation (Extended Data Figure 5). As before, the majority of ribosomes were stalled with a peptidyl-tRNA in the P site (Extended Data Figure 7). The map of the stalled MYC complex without IDB-001 present reached a resolution of 2.2 Å (2 Å local resolution in the PTC), and the stalled MYC:IDB-001 complex reached a resolution of 1.9 Å (1.8 Å local resolution in the PTC) (Extended Data Figure 8). As in the previous structures, the identity of the nascent chain in positions 0 to –3 could be well-modeled in both maps (Figure 4A-B). Because the two structures only differ from the presence/absence of IDB-001, we could directly compare the conformational changes resulting from compound binding. An overlay of the two structures showed that the general backbone of the polypeptide was in a similar position in the PTC; however, a key difference was the change in the orientation of the aspartic acid at position –1 (4-mer D_-1_, MYC D_258_) (Figure 4C). In the absence of IDB-001, the density for the side chain D_-1_ is oriented down and away from the compound binding pocket (Figure 4A). In contrast, in the presence of IDB-001, D_-1_ is clearly oriented toward the distal pyrrolidine ring, resulting in a stabilizing salt-bridge, similar to the earlier observation made for IDB-001 in the presence of the FTED peptide (Figure 4B-C). This suggests that structural rearrangements within the peptide chain can be achieved by a complimentary drug molecule to form new high affinity interactions.

**Figure 4.**
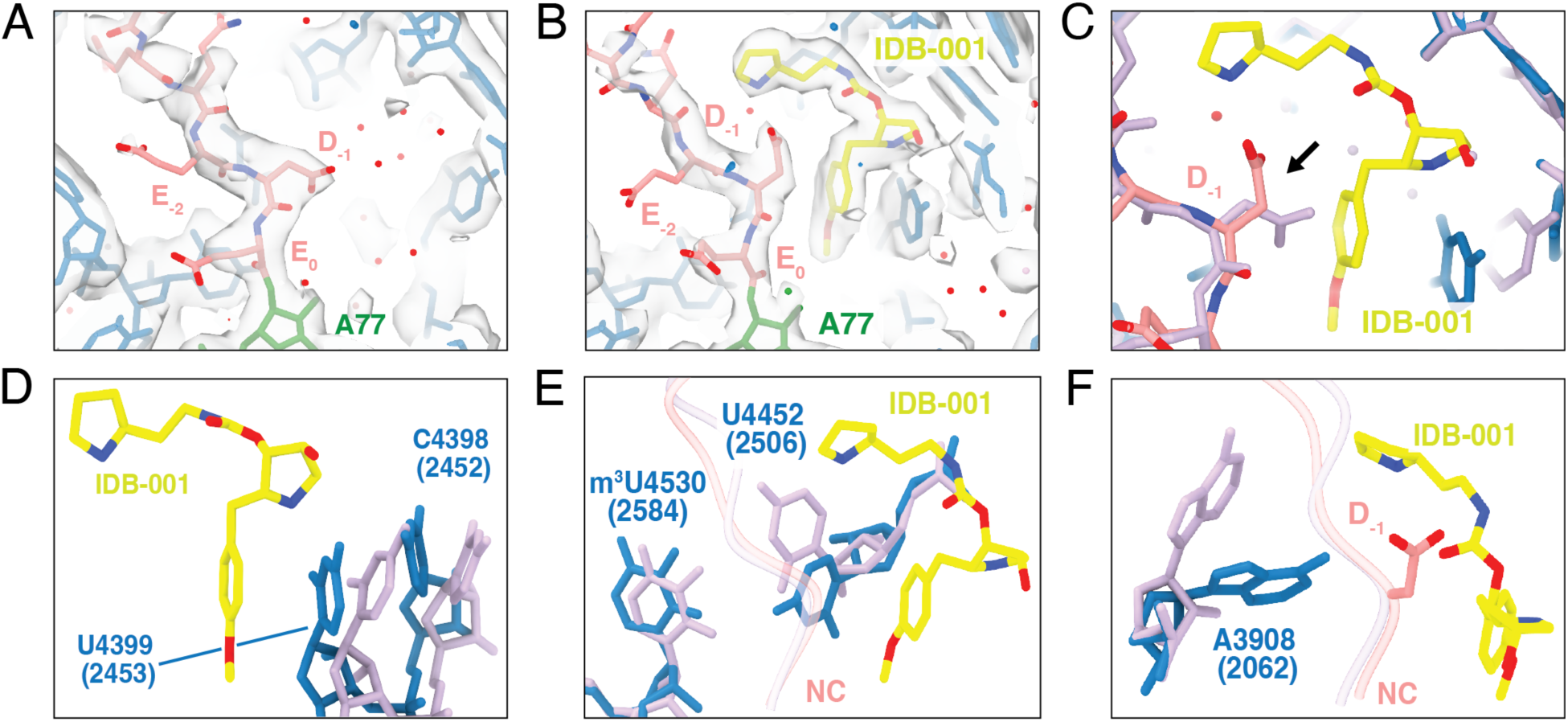
Structural rearrangements upon binding of IDB-001 to a MYC peptide-containing RNC. A. Closeup of the PTC for the MYC-containing RNC structure in the absence of any interdictor. The nascent MYC peptide Q_-3_E_-2_D_-1_E_0_ is captured in the PTC attached to the P-site tRNA. B. Same as A, but in the presence of IDB-001. C. Overlay of the atomic models from A and B, showing the rearrangement of the of Asp (D_-1_) side chain upon IDB-001 binding (arrow). rRNA nucleotides from the apo (minus IDB-001) structure are colored lavender, while nucleotides from the +IDB-001 are colored blue. D-F. Conformational changes of the rRNA induced by the presence of IDB-001 with focus on C4398, C4399 (D), U4452, U4530 (E) and A3908 (F). Colors as indicated in (C).

Comparison of the cryo-EM structures with and without bound interdictor also allowed us to determine conformational changes induced by the compound on rRNA in the PTC. Similar to observations with ANS^2,30^, we find that binding of IDB-001 to the ribosome causes the bases of residues C4398 and C4399 (*E.c. 2452 and* 2453) to move by several angstrom towards the compound to engage in π-stacking interactions, compacting the space in the PTC (Figure 4D). Moreover, several other bases implicated in the peptidyl-transfer reaction change their orientations in the presence of compound. In the absence of compound, U4452 (*E.c.* 2506), which acts as a gate to control access of the A-site aminoacyl-tRNA to the sensitive P-site tRNA-peptidyl ester bond, alternates between two conformations separated by a 90-degree swing^42–45^. In the presence of IDB-001, U4452 is instead forced into a position that leaves the gate half closed, potentially affecting peptidyl-transfer (Figure 4E). A3908 (*E.c.* 2062), which lines the wall of the exit tunnel adjacent to the PTC, has also been shown in previous structures to adopt two distinct conformations that could interact with the growing polypeptide chain^15^. It has been hypothesized that molecules bound in the A site could shift the equilibrium between these two states, impacting nascent chain movement^15,16,46^. In the presence of IDB-001 the equilibrium of A3908 is shifted towards the ‘bottom’ conformer, closer to the compound, potentially hydrogen bonding to the repositioned sidechain of the –1 aspartic acid (Figure 4F). Together, our comparative analysis shows that interdictors cause distinct changes in the PTC that allow them to bind and specifically engage with the nascent chain. Taken together, our results and previous insights from earlier mechanistic studies of ANS^2,30^, lead us to conclude that interdictors are tri-functional: the core pyrrolidine affords binding affinity to the PTC of the ribosome while the distal R groups confer differential biochemical selectivity to the nascent chain. On the other end of the molecule, the methoxyphenyl group likely impedes accommodation and transpeptidation of incoming A-site tRNAs.

### Interdictors reduce cancer cell viability and differentially regulate the ribotoxic stress response

We next sought to confirm the potential for these selective small molecule ribosome stallers to impact cancer cell fitness by performing cell viability assays in a small set of MYC-dependent cancer cell lines^47^ including HCC-1143 (TNBC), 22Rv1 (prostate cancer), LS411N (colorectal cancer), and MCF7 (HR+, HER2-breast cancer) (Figure 5A & Table 1). Both compounds reduced cancer cell viability, though IDB-002 was significantly more potent in both LS411N and 22Rv1 cells and similar in potency to ANS. Given the MYC-dependent nature of these cell lines, and that the MYC protein has a very short half-life in highly proliferative cells, we asked if treatment with interdictors would lead to the depletion of MYC. Indeed, Western blot analysis confirmed rapid MYC reduction (< 4 hours) in HCC-1143 cells (Figure 5B), consistent with MYC’s short half-life in these cancer cells (Extended Data Figure 9A).

**Figure 5.**
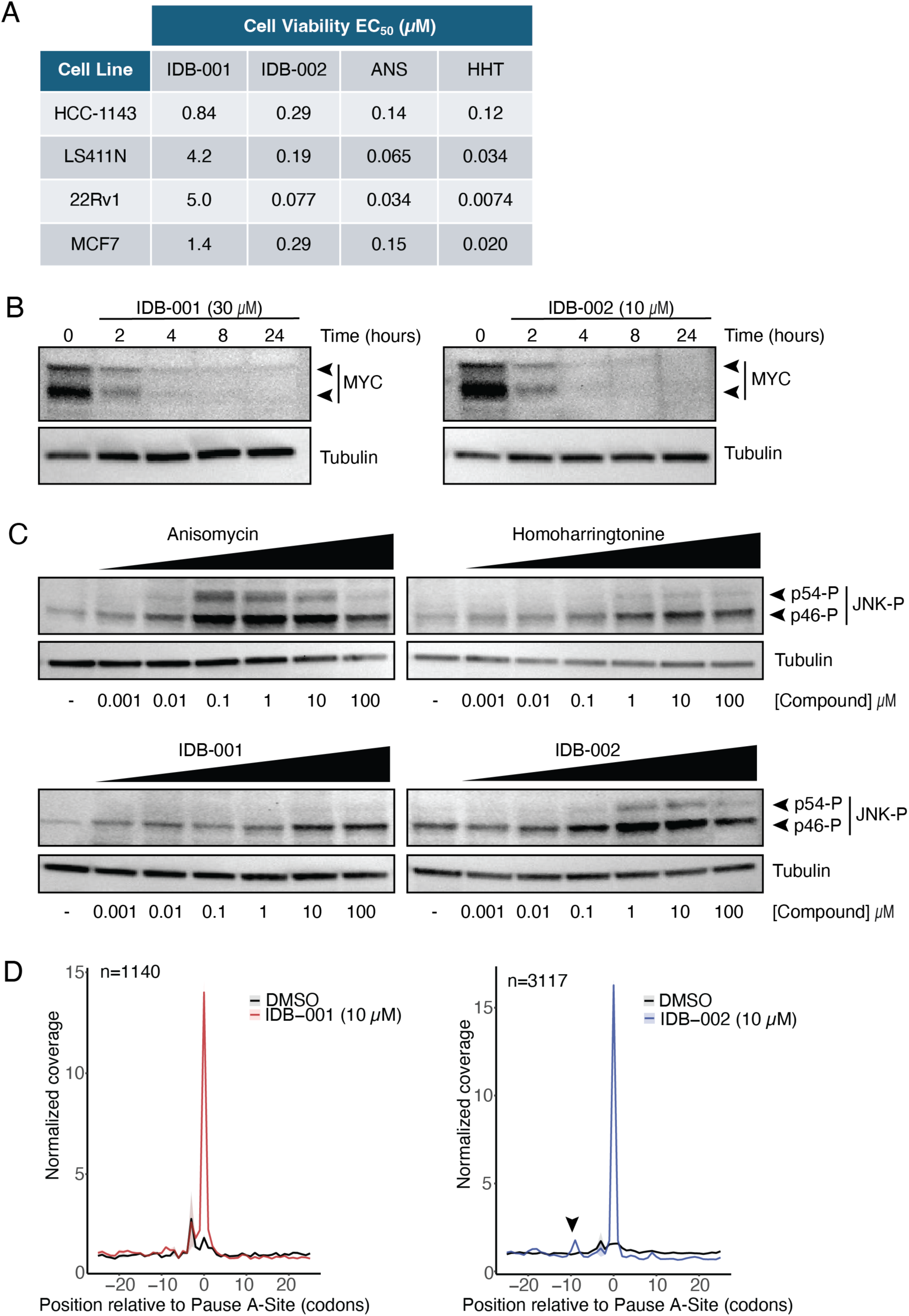
Interdictors inhibit MYC-dependent cancer cell viability and lead to MYC depletion. A. Cell viability EC_50_ measurements for IDB-001, IDB-002, anisomycin (ANS) and homoharringtonine (HHT) across a panel of MYC-dependent cancer cell lines. Data including confidence intervals from curve fitting are included in Table 1. B. Western blot analysis of MYC protein in HCC-1143 triple-negative breast cancer cells treated with either IDB-001 or IDB-002 at 30x the IC_50_ concentration (30 µM for IDB-001, and 10 µM for IDB-002). C. Western blot analysis for JNK phosphorylation (JNK-P), a marker of ribotoxic stress pathway with varying interdictor or translation inhibitor treatment concentrations in HCC-1143 cells. D. Metagene analysis of ribosome occupancy in the regions flanking pause sites for IDB-001 and IDB-002. Pause sites filtered to exclude regions within 25 codons of start codon and overlapping sites. Arrowhead indicates peak suggestive of ribosome collision upstream of pause site.

**Table 1.** 72-hour cell viability assay EC_50_ measurements with 95% confidence interval.

| **Cell Line** | **Compound** | **EC<sub>50</sub> (μM)** | **EC<sub>50</sub> Upper CI (μM)** | **EC<sub>50</sub> Lower CI (μM)** |
| --- | --- | --- | --- | --- |
| 22RV1 | IDB-001 | 4.983 | 6.649 | 3.735 |
|  | IDB-002 | 0.077 | 0.103 | 0.058 |
|  | IDB-003 | 0.042 | 0.053 | 0.033 |
|  | ANS | 0.034 | 0.045 | 0.026 |
|  | HHT | 0.007 | 0.010 | 0.006 |
| HCC1143 | IDB-001 | 0.837 | 1.152 | 0.608 |
|  | IDB-002 | 0.288 | 0.362 | 0.229 |
|  | IDB-003 | 0.132 | 0.152 | 0.115 |
|  | ANS | 0.143 | 0.177 | 0.115 |
|  | HHT | 0.124 | 0.149 | 0.104 |
| LS411N | IDB-001 | 4.233 | 5.098 | 3.514 |
|  | IDB-002 | 0.191 | 0.236 | 0.154 |
|  | IDB-003 | 0.047 | 0.053 | 0.041 |
|  | ANS | 0.065 | 0.084 | 0.051 |
|  | HHT | 0.034 | 0.042 | 0.028 |
| MCF7 | IDB-001 | 1.364 | 1.648 | 1.129 |
|  | IDB-002 | 0.295 | 0.341 | 0.255 |
|  | IDB-003 | 0.063 | 0.070 | 0.056 |
|  | ANS | 0.149 | 0.206 | 0.108 |
|  | HHT | 0.020 | 0.026 | 0.016 |

Given their role in inhibiting translation elongation, we asked whether these compounds would stimulate the ribotoxic stress response (RSR) as has been documented for ANS in eukaryotic cells^27,48–50^. Intermediate concentrations of ANS which lead to high frequency ribosome collisions stimulate the RSR and trigger JNK-mediated apoptotic signaling^27^. In HEK293T cells, we reproduced the reported concentration-dependent, robust increase in phosphorylated JNK caused by ANS (Extended Data Figure 9B). We also observed this increase for IDB-002 but saw no phosphorylated-JNK on treatment with either IDB-001 or HHT. In HCC-1143 cells, we observed a similar trend, but with the distinction that both ANS and IDB-002 led to a less pronounced increase in JNK-P levels compared to that observed in HEK293T cells (Figure 5C & Extended Data Figure 9B). Additionally, IDB-001 and HHT led to a subtle increase in phosphorylation of the p46 isoform of JNK at concentrations above 1 μM, while leaving the p54 isoform unchanged (Figure 5C). In both cell lines, RSR activation by IDB-002 was higher than with IDB-001, suggesting that IDB-002 might cause more ribosome collisions than IDB-001. Indeed, when we perform a global metagene analysis of the ribosome footprints centered around pauses, we observe a stronger peak corresponding to ribosome pileup with a drop-off of density after the pause site for IDB-002 treatment (Figure 5D). These data confirm that interdictors can inhibit the viability of therapeutically relevant cancer cell lines either concomitant with, or independent of the RSR.

### IDB-003 inhibits tumor growth of MDA-MB-231 xenografts *in vivo*

We next wanted to establish *in vivo* proof-of-concept for this modality for inhibition of xenograft tumor growth in mice. To further enhance drug-like properties and enable oral dosing for an advanced tool molecule, we replaced the methyl ether in IDB-002 with an oxazole, resulting in the *in vivo* ready compound IDB-003 (Figure 6A). *In vitro*, IDB-003 displayed modestly improved potency compared to IDB-002 in the set of MYC-dependent cancer cell lines (Figure 6B) and similarly led to rapid MYC depletion in HCC-1143 cells (Extended Data Figure 10A). Additionally, the shared northern substituent of IDB-002 and IDB-003 that interacts with the nascent polypeptide in the PTC imparts a similar pattern of context-dependent activity observed by ribosome profiling experiments (Figure 1G & Extended Data Figure 10B). To test the efficacy of IDB-003 on solid tumor growth inhibition *in vivo*, we chose the highly metastatic and difficult to treat TNBC model MDA-MB-231^51,52^, which has been shown to be less-sensitive to FDA-approved CDK4/6 inhibitors because of robust MYC expression^53^. After 28 days of twice daily oral dosing (BID p.o.), we observed 80% tumor growth inhibition (TGI) for mice given a 10 mg/kg dose of IDB-003, and 58% TGI for the group given a 3 mg/kg dose compared to vehicle control. These data strongly support that interdiction of protein synthesis is an efficacious and viable strategy for the treatment of solid tumors *in vivo*.

**Figure 6.**
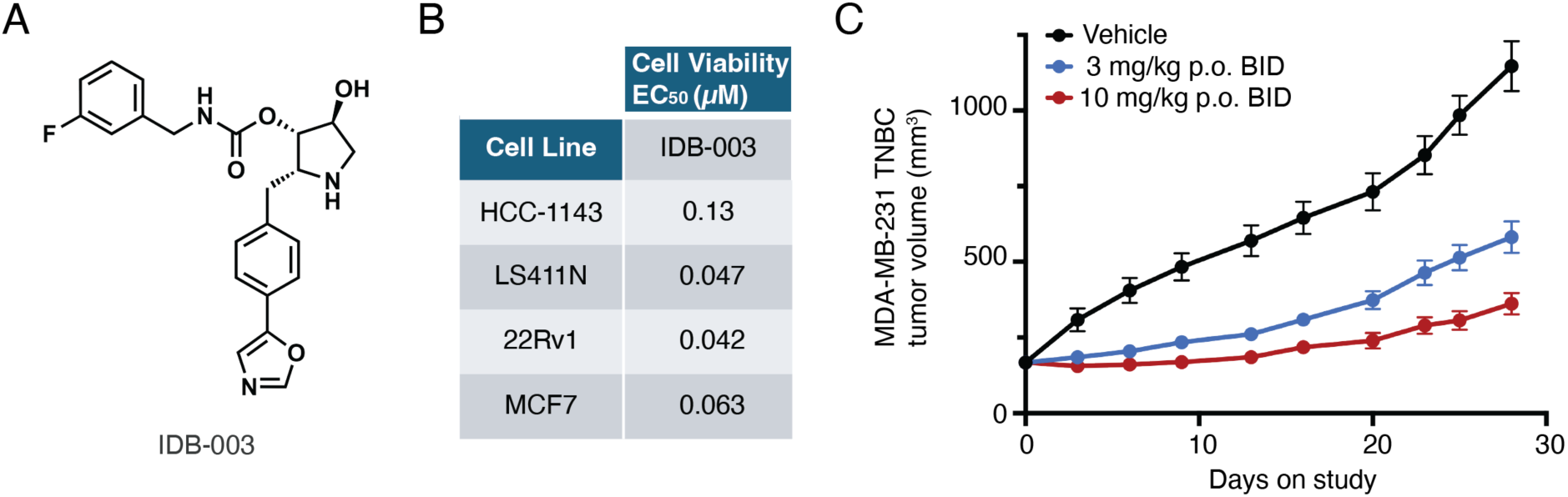
IDB-003 inhibits MDA-MB-231 tumor growth *in vivo*. A. Chemical structure of interdictor IDB-003 which has improved drug-like properties compared to IDB-002 and is used in the *in vivo* efficacy study. B. Cell viability EC_50_ measurements for IDB-003 across a panel of MYC-dependent cancer cell lines. Data including confidence intervals from curve fitting are included in Table 1. C. Tumor volume measurements recorded throughout the 28-day efficacy study of mice bearing MDA-MB-231 xenograft tumors and treated with indicated doses of IDB-003 (n=8 per group) or vehicle (n=6). Error bars are SEM.

## DISCUSSION

In this report, we demonstrate the ability to engineer the sequence-selective action of PTC inhibitors of the human ribosome and showed that designed analogs can inhibit MYC-driven cancer cell proliferation. In two representative, structurally distinct compound sub-series, we report context-dependent ribosome stalling both *in vitro* and *in cellulo* that we can attribute to the physicochemical-selective actions induced by specific chemical moieties appended on a shared scaffold to interact with the nascent polypeptide in the PTC. IDB-001, with a basic substituent, showed preference for complementarily negatively charged residues, while IDB-002 with a non-polar substituent showed preference for hydrophobic residues (Figures 1-2). Moreover, our ribosome profiling allowed us to document for the first time, the sequence preferences for ANS, a natural product inhibitor of translation elongation canonically thought to be an indiscriminate elongation inhibitor (Figure 1E, Extended Data Figure 1D)^54,55^.

The sequence-selective stalling we observe for the reported molecules is primarily mediated through interactions with residues at the penultimate (–1) position of the nascent chain (Figure 1F-G), although we also identify distinct contributions from the neighboring amino acids upon compound treatment (Figure 1I-J). This degree of context-dependent activity is reflected in our observations of concentration dependence, with a shift in global ribosome occupancy towards the 5’ end of coding sequences (Figure 1B) at high concentrations, while at lower concentrations (Extended Data Figure 2) we observe a much less significant polarity shift with correspondingly fewer stall events induced by the compounds (Figure 2A-B). Ongoing chemical library expansion guided by these preliminary findings with these early tool molecules is expected to lead to molecules which are increasingly more selective for a broader collection of particular sequence motifs.

In addition to the stalling mediated through engagement of a given peptide sequence, we found that relative position within the coding sequence influenced the likelihood of interdiction. Specifically, ribosomes decoding amino acids 5-10 of any given ORF preferentially exhibit stall signatures (Extended Data Figure 1A). This location is particularly intriguing to us as it represents the region in which a trailing ribosome would not be able to initiate on the mRNA due to steric constraints^56^. These positions therefore represent sites in which a single ribosome can be stalled without potentially causing a ribosome-ribosome collision, which are known to trigger both mRNA and ribosome quality control pathways^48,49,57,58^. While we can observe this at the metagene level, it is difficult to decipher whether this phenotype derives from the compound’s preferential binding, or these stalled monosomes are simply enriched in our profiling libraries due to an effect in stabilizing these mRNAs. Further exploration into the positional dependency and contribution of local sequence impacts, including codon optimality, intrinsically slow regions, and RNA structure^3^ will inform continued exploration of the nuances of this novel therapeutic modality as well as target identification for rational design.

On a molecular level, our structural studies reveal that interdictors bind in a linkerless heterobifunctional manner to both the rRNA and the nascent peptide chain, where each independent binding interaction can largely be attributed to separate parts of the comparatively compact small molecule framework (Figures 3-4 & Extended Data Figure 6). As these functions are somewhat decoupled from each other, though still operate in concert, we have shown that they can be independently modulated in ongoing structure-based design efforts, much the way a PROTAC^59,60^ can be modularly designed and constructed. In particular, the functional substituent projecting towards the polypeptide chain can be further modified to leverage shape and charge complementarity and to enable interaction with the nascent chain beyond the residues from the 0 to the –3 position described herein. Similar considerations also explain the observed de-enrichment of certain sequences that contain bulky aromatic residues, as they are likely to clash sterically with the molecule in the PTC (Figure 1I-J). Furthermore, A site occlusion can result from the positional anchoring of the molecule induced by these binding contacts. Together, modifications to each of these functions individually, or in combination, will continue to contribute to the expansion of a diverse and efficacious library of compounds which selectively target various peptide motifs in the PTC.

Herein we also report >15-fold selectivity for different analogues to kinetically inhibit translation elongation of stall motif-containing mRNAs (Figure 2E-F), highlighting a baseline biochemical selectivity that may be achieved by this modality. This basal biochemical selectivity is subsequently magnified by target-intrinsic properties such as half-life/resynthesis rate, adding a very important temporal aspect to drug action. The concept of sequence selectivity of small molecule inhibitors of translation may be thematically similar to the sequence selectivity of RNA interfering therapies that also target mRNA translation; however, orally dosed PTC-binding molecules circumvent the significant challenges of delivery and stability of short RNAs^61^. Furthermore, unlike siRNAs or small-molecule mRNA binders that inhibit translation by binding to mRNA directly^62^, PTC-binding molecules benefit from leveraging the diversity of molecular interactions that can be made with twenty amino acid side chains rather than only four nucleobases, and, similarly to targeted protein degradation approaches, benefit from a sub-stoichiometric or catalytic mode of action.

Because interdiction is agnostic to the three-dimensional structure of target proteins outside the ribosome, this approach is uniquely positioned to address diseases dependent on traditionally “undruggable” target genes that do not have structured binding pockets. We, as others^24,25^, hypothesize that translation inhibition would be especially effective for targeting oncogenes with short intrinsic half-life or fast turnover^63^ like MYC, while longer-lived housekeeping genes^64^ will not be appreciably impacted, adding an additional layer of pharmacological selectivity. Indeed, we confirm that all three tool interdictors presented here lead to a rapid depletion of MYC protein from cells to inhibit their viability (Figures 5-6 & Extended Data Figure 10). Furthermore, a more advanced tool molecule, IDB-003, is highly effective *in vivo* at low, well-tolerated oral doses in the difficult to treat MDA-MB-231 TNBC model (Figure 6C). Together, these data validate our approach of sequence-selective interdiction of protein synthesis as a powerful therapeutic strategy for MYC-driven cancers, and more completely optimized molecules are progressing into pre-clinical development.

We fully expect that this approach of engineered protein target sequence-selectivity can be extended to many other short half-life oncogenes and transcription factors beyond MYC. Finally, while the inherent turnover rate of a target for interdiction can be a particularly significant factor in oncology settings, where target deprivation to an addicted cancer cell can drive downstream cell viability within a relatively short window, it does not limit the application of this modality to oncology. We expect that the universal nature of the modality can be extended to many protein factors critical in neurodegeneration, immunology, and other therapeutic areas where even a more modest reduction in translation could have a significant positive impact on disease progression.

## MATERIALS AND METHODS

### Cell culture, drugging, and harvesting for ribosome profiling

HCC1143 breast cancer cells (ATCC) were cultured in RPMI 1640 medium with 10% fetal bovine serum (FBS), 100 U/mL penicillin, 100 U/mL streptomycin and 1% GlutaMax (Gibco). For ribosome profiling experiments, media was replenished 24 h prior to harvesting. When cells reached 70-80% confluency, they were treated with anisomycin (ANS) or interdictor for 60 min at 10 µM (unless otherwise noted). Cells were washed with ice-cold phosphate-buffered saline and lysed in lysis buffer (20 mM Tris-HCl pH 7.4, 150 mM NaCl, 5 mM MgCl_2_, 1 mM DTT, 1% v/v Triton X-100, 25 U/mL Turbo DNase I). Lysates were incubated on ice for 30 min to allow for DNase digestion and clarified by centrifugation at 20,000 × g for 30 min at 4 °C. Lysate RNA content was quantified with Quant-it™ RiboGreen reagent (Thermo Fisher) on an EnVision microplate reader (Revvity), following the manufacturer’s instructions. Cycloheximide was not used in this study.

### Ribosome profiling library preparation

Lysates were digested with 3.33 U/µg RNase I (Ambion) for 45 min at room temperature with gentle agitation. Monosomes were then isolated by sucrose cushion, using 450 µL 1M sucrose in polysome buffer (20 mM Tris-HCl pH 7.4, 150 mM NaCl, 5 mM MgCl_2_, 1 mM DTT) for 400 µL of digested lysate material. Sucrose cushions were performed with MLA-130 rotor in an Optima Max-XP ultracentrifuge (Beckman Coulter) at 100,000 × g for 90 min at 4 °C. Pellets were resuspended in resuspension buffer (150 mM NaCl, 20 mM EDTA, 1% Triton X-100, 20 mM pH 7.5 Tris-HCl) and incubated at room temperature for at least 10 min. RNA was extracted using Direct-zol RNA miniprep kit (Zymo), according to the manufacturer’s instructions. Ribosome footprint RNA size selection was performed as previously described^65^ with minor modifications, including the use of 10% TBE-urea gels (Biorad) and custom sizing markers (oIB188-190, 17nt-34nt). Following overnight gel extraction and ethanol precipitation, RNA was dephosphorylated with T4 PNK (NEB) at 37 ° C for 1 h. Ribosome footprint cDNA libraries were subsequently constructed using Ordered Two-Template Relay kits (Karnateq) according to the manufacturer’s recommendations^32^. cDNA libraries were indexed and amplified with NEBNext Indexing Primers (NEB).

### Ribosome profiling data processing

Raw ribosome profiling sequencing reads were first trimmed to remove adapter sequences using Cutadapt^66^, discarding any reads shorter than 16 nucleotides. The trimmed reads were then mapped sequentially to rRNA, cytoplasmic and mitochondrial tRNA references, followed by alignment to mitochondrial mRNA, mitochondrial DNA, and various noncoding RNA references with Bowtie and Bowtie2^67,68^. Next, any remaining reads were aligned to human mRNA (RefSeq MANE v1.3) and processed through RSEM^69^ for transcript quantification. RSEM quantifications of mRNA-aligned RFPs were used for all downstream analyses. Using the length of each ribosome footprint and the reading frame identified at the 5’ end of the ribosome-protected fragment, the A-site codon was inferred and subsequently used to determine the corresponding peptide sequence in the nascent polypeptide chain.

### Pause site identification and sequence analysis

To identify codon positions at which ribosome occupancy increased in response to compound, we first restricted our analysis to genes with an average coverage exceeding 0.5 reads per codon across replicates. We then adapted a previously described strategy^36,70^, employing two-tailed Fisher’s exact tests to identify codons displaying significant changes in ribosome occupancy between compound and DMSO-treated samples. Briefly, read counts at each codon were average across replicates, rounded to nearest integer, used to generate 2×2 contingency table comparing the ratio of compound-versus control-derived ribosome footprints at a given codon against the ratio of all other codons in the same gene. Multiple-testing correction was performed using the Bonferroni method. A codon was deemed a pause site if it had an odds ratio greater than 1 (and finite) and its adjusted p-value was less than 0.05. Finally, we used ggseqlogo^37^ to create sequence logos of these pause sites (from positions –7 to 0) by scaling the nascent peptide sequences of the pause sites by the corresponding odds ratio.

### Metacoverage and polarity scoring analyses

To examine how interdictor treatment influenced the distribution of ribosome footprints along the length of transcripts, we carried out metacoverage and polarity scoring analyses. First, each ribosome footprint was assigned to the 5′ UTR, the coding sequence (CDS), or the 3′ UTR. A normalized meta-position value was calculated by dividing the footprint’s location by the length of its respective region in that transcript, then shifting these values so that entire transcript could be represented from meta-position –1 to +2 with: −1 to 0 corresponding to 5′UTR, 0 to +1 representing CDS, and +1 to +2 representing 3′ UTR. Only footprints from genes with adequate coverage (>0.1 reads per codon) were retained. At each meta-position (rounded to two decimal places), counts were normalized to library size and averaged across replicates.

We next calculated a polarity score for each transcript (adapted from^34^), which captures the distribution of ribosome coverage from the start to the stop codon. Specifically, for a CDS of length *L* codons, the polarity score P is given by:

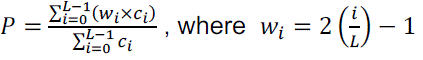

Here, *c_i_*_’_ is the ribosome footprint coverage (read count) at codon *i*, and *w_i_*_’_ is a position-specific weight ranging from –1 at the start codon (*i* = 0) to +1 at the stop codon (*i* = *L*). Thus, P varies between –1 (indicating that all ribosomes were positioned at the start) to +1 (indicating that they were entirely clustered at the stop codon).

### Differential ribosome occupancy analysis

Ribosome footprints and corresponding nascent peptide sequences spanning four codons (positions –3 to 0) were used to generate count tables corresponding to the number of reads mapping to each unique 4-mer across samples. To identify 4-mers with changing ribosome occupancy in compound-treated versus DMSO-treated samples, differential expression analysis across 4-mers was performed with DESeq2^38^. 4-mers showing significant increase in ribosome occupancy (log_2_ fold-change > 0.5) were scaled according to their fold-change and input into ggseqlogo^37^ to generate peptide motif logo.

Similarly, the position-weighted matrix analysis was performed by extracting amino acid identities from ribosome footprints and corresponding nascent peptide sequences from nascent chain position (–7 to 0). After filtering for coverage threshold (genes with > 0.1 read/codon) and normalizing to library size, log_2_ fold-change was calculated by comparing each treatment condition to DMSO for every residue at each position.

### Selection of 4-mer sequences for the *in vitro* luminescence assay

To ensure that the majority of the signal from interdiction of the 4-mer containing reporters was derived from interdiction of the 4-mer sequence rather than the HiBit tag we tiled six repeats of each 4-mer before the 11 AA HiBit tag. Due to tiling of the 4-mer sequence in this way the ribosome will encounter, and present for interdiction, not just the 4-mer of interest but also all four ‘rotations’ of that 4-mer. Therefore, we created a ‘rotation score’ metric that summed the log2-fold change for all four ‘rotations’ of the 4-mer, and chose 4-mers for inclusion in the luminescence assay based on this score. As such the expected level of interdiction, estimated from the ribosome profiling experiments, is based on the entire 24 AA reporter sequence rather than the single 4-mer ‘hit’.

### Creation of reporter plasmids for the *in vitro* transcription-translation luminescence assay

The plasmid pT7CFE1 (Thermo Scientific) was digested in rCutsmart buffer with the restriction enzymes XhoI and MscI (New England Biolabs) for 1 h at 37 °C. The linearized plasmid was used as a template for Gibson Assembly together with a gBlock (Integrated DNA Technologies) containing the coding sequence of interest and the HiBit tag, followed by a stop codon. Gibson assembly was carried out using a Gibson master mix (New England Biolabs) at 50 °C for 1 h. 2 µL of the Gibson assembly reaction was used to transform 20 µL of Mach 1 competent E. coli (Invitrogen). The correct plasmid sequence was verified by whole plasmid sequencing (Genewiz).

### *In vitro* transcription-translation luminescence reporter assay

An *in vitro* translation transcription system was created by combining on ice, for each sample to be measured, 5 µL of HeLa cell lysate (Ipracell), 1 µL of Accessory protein mix (Thermo Fisher), 2 µL of Reaction mix (Thermo Fisher), and 1 µL of HiBit substrate mix, consisting of 1:10 Nanoluc substrate (Promega) and 1:10 LgBit protein (Promega) in Nanoluc sample buffer (Promega). To this mixture was added 2 µL of interdictor dissolved in water at the required concentration. The reaction was started by addition of 2 µL of the relevant reporter plasmid at 110 ng/µL and transfer of the assay mixture to 30 °C within a luminescence-capable plate reader (BMG Labtech). The ensuing *in vitro* transcription-translation reaction was then followed in real time by recording the luminescence intensity of each sample every 108 s. The resulting luminescence time traces were then analyzed in Prism 10 (GraphPad Software) by identifying the linear portion of each trace, corresponding to steady-state protein synthesis, and determining its slope through linear fitting. The slopes of each reaction where then plotted against the interdictor concentration and IC_50_ values determined by non-linear fitting of the equation “Inhibitor vs. Response – three parameters” within Prism 10.

### Generation of stalled RNC complexes for cryo-EM

Stalled RNC complexes for cryo-EM were generated by *in vitro* translation of nonstop mRNAs as previously described^41^. Linear DNA template encoding an IRES, followed by a 3x FLAG tag, the Villin head domain^71^ and the sequence of interest was amplified by PCR using a reverse primer that contained two extra nucleotides carrying a C2’-methoxy modification proximal to the 5’ end to reduce non-templated additions in the subsequent T7 transcription step^72^. Linear DNA was added to IVTT reactions (Thermo Scientific 1-Step Human Coupled IVT Kit, 88881) according to manufacturer’s instructions and incubated with HeLa cell lysate for 20 min at 30 °C. Reactions were quenched with 1 mM buffered cycloheximide and clarified by centrifugation for 20 min at 4 °C and 20,000 rcf. Reactions were pelleted through a 1 M sucrose cushion containing 750 mM KOAc, 5 mM MgOAc and 20 mM Hepes pH 7.4 at 100,000 rpm for 2 h at 4 °C in a TLA100.3 rotor (Beckman Coulter). Reactions were resuspended in 20 mM Tris pH 7.5, 100 mM KOAc, 7.5 mM MgOAc, 0.25 mM spermidine, and 0.01 % NP-40 and incubated with pre-equilibrated M2 FLAG bead slurry (Pierce) at 4 °C for one h. Beads were washed with resuspension buffer, including a high salt wash step with KOAc adjusted to 750 mM before elution with 1 mg/mL 3X FLAG peptide (Sigma). Beads were regenerated with 0.1 M glycine and the re-incubated with the flow-through from the previous step. The eluates were combined and concentrated in a 100 kDa cutoff centrifugal spin filter (Amicon) to remove FLAG peptide, smaller contaminants and to exchange the buffer to cryo-buffer [20 mM Tris pH 7.5, 100 mM KOAc, 7.5 mM MgOAc, 0.25 mM spermidine, 0.05 % Octyl-β-D-glucopyranoside (Thermo Scientific)].

### Cryo-EM grid preparation, data collection and processing

Between 150 and 300 nM of purified RNC complexes were incubated on ice with 50 µM of compound that has been pre-diluted in cryo buffer for 30 min. 4 µL of sample were applied on glow discharged Cu 2/1 300 mesh + 2 nm carbon grids (Quantifoil) and plunge frozen in liquid ethane using a Vitrobot (Thermo Scientific), using a blot force of 0 and a blot time of 1 s. Grids were imaged a the LSI Cryo-Electron Microscopy Lab at the University of Michigan. Data was collected on a Titan Krios microscope (Thermo Scientific) operating at 300 kV and equipped with a Gatan K3 direct electron detector and a Bioquantum imaging filter. Movies were acquired using a pixel size of 0.831 Å/px, a total dose of 50 e^-^/Å^2^ and using a nominal defocus range between – 0.8 µm to –1.5 µm. All data processing was carried out in Cryosparc v4.5^73^ and is schematically outlined in Extended Data Figure 7. After motion correction and CTF estimation, particles were picked using the blob picker with a particle size of 250-300 Å. After particle extraction and 2D classification, ribosome classes were refined to a consensus structure that was used as a basis for 3D classification without alignment using a filter of 12 Å to identify conformationally and compositionally distinct ribosome classes. Repeated rounds of 3D refinement and 3D classification without alignment with focus on each tRNA yielded the final classes that are shown in Extended Data Figure 7. Classes of interest were refined to high resolution by performing per-particle CTF correction and reference-based motion correction, followed by non-uniform refinement and local refinement for regions of interest. Resolution of each reconstruction was assessed using the FSC = 0.143 criterion^74^.

Final, unsharpened maps were used for atomic modelling in Isolde v1.8^75^ and Coot v1.1.1^76^, using PDB 8GLP as a starting model^30^. B-factor sharpened maps were used for placing water molecules, ions and ligands. Final macromolecular refinement was carried out in Phenix v1.21.2^77^, using restraints generated in elBOW for modified nucleotides and compounds. Maps and models were visualized and analyzed and structure figures created with ChimeraX v.1.8^78^.

### Cell Culture and Western Blot analysis

All cell lines used in this study were purchased from ATCC and cultured using recommended protocols. Western blot analyses were performed by plating 5 million cells in 15 cm dishes with the appropriate media and leaving cells to adhere overnight. The next day, compounds were diluted in 100% DMSO and directly added to each dish, with a final concentration of 0.5% DMSO or less. Sample timepoints were rinsed with 1 mL of cold PBS at time of collection, scraped and pelleted (5 min at 500 x g). The PBS was then aspirated off and cell pellets frozen in LN_2_. Frozen cell pellets were lysed on ice for 20 min in 50 µL of RIPA buffer (Thermo Scientific, 89901) supplemented with HALT Protease Inhibitor with EDTA (Thermo Scientific, 78429), HALT Phosphatase Inhibitor (Thermo Scientific, 78420), and benzonase (Sigma-Aldrich, E1014-25KU). Lysates were clarified by centrifugation at 13,000 rpm for 10 min. After clarification, supernatants were quantified by BCA analysis (Thermo Scientific, 23225) or equivalent loading and denatured in SDS-PAGE dye. Proteins were resolved by 4%–15% Criterion TGX midi protein gels (Bio-Rad, 5671084) and transferred to PVDF membranes using a Trans-Blot Turbo transfer system (Bio-Rad). Non-phosphorylated (c-MYC, Tubulin) membranes were blocked with 5% nonfat dry milk (Cell Signaling, 9999S) in TBST with primary antibody overnight with gentle rocking at 4°C, while phosphorylated (P-JNK) membranes were treated similarly except for using 5% BSA in TBST. After overnight incubations, blots were washed three times for 5 min in TBST and then incubated with secondary antibodies at RT with gentle rocking for 2 h. Membranes were washed again with TBST before being developed using SuperSignal West Femto Maximum Sensitivity Substrate (Thermo Scientific, 34096) and imaged on an iBright system (Invitrogen).

### Cell Viability Assay

Indicated cell lines were plated in black Bio-One CELLSTAR 96-well, cell culture-treated, flat-bottom microplates (Greiner) at concentrations between 2,000-10,000 cells/well in tissue culture media and incubated overnight in standard tissue culture incubator (37°C, 5% CO_2_). Once cells adhered, compounds resuspended in 10% DMSO were added to each well containing 100 uL of media for a final concentration of 0.5% DMSO. After 72 h of incubation, viability was assessed using established protocols for CellTiter-Fluor assay (Promega). Briefly, the media was removed from the plates, cells gently rinsed with PBS, and a 1:1 mix of PBS and fluorescent substrate is added to the plates before incubating for 60 min at room temperature with orbital shaking. Finally, viability was quantified per kit instructions using an Envision fluorescent plate reader (Revvity). To calculate EC_50_ values, each plate is background subtracted and normalized to DMSO only control wells for a % viability compared to control. Non-linear regressions for each compound and cell line were performed to obtain the EC_50_ value.

### MDA-MB-231 xenograft study in mice

Efficacy study for IDB-003 in human TNBC MDA-MB-231 tumor bearing, female BALB/c nude mice was performed at Pharmaron (Beijing, China). All procedures related to animal handling, care, and treatment in this study were performed according to guidelines approved by the Institutional Animal Care and Use Committee (IACUC) of Pharmaron and in compliance with the guidance of the Association for Assessment and Accreditation of Laboratory Animal Care (AAALAC). In addition, all portions of this study were performed at Pharmaron and adhered to the study protocol provided or approved by the sponsor and applicable standard operating procedures (SOPs). Animals were quarantined for 7 days before the start of any experimental procedure. The general health of the animals was evaluated by a veterinarian, and complete health checks were performed. Animals with abnormalities were excluded prior to the study start. General procedures for animal care and housing were in accordance with the Commission on Life Sciences, National Research Council, and standard operating procedures (SOPs) of Pharmaron, Inc. The mice were kept in laminar flow rooms at constant temperature and humidity with 4-5 animals in each cage. Animals were housed in polycarbonate cage which was the size of 300 × 180 × 150 mm^3^ and in an environmentally monitored, well-ventilated room maintained at a temperature of 23 ± 3°C and a relative humidity of 40% 70%. Fluorescent lighting provided illumination approximately 12 h per day, lights on 8 AM, and off at 8 PM. Animals had free access to irradiation sterilized dry granule food during the entire study period, except where specified in the protocol. Hydrogel supplementation were offered to animals if and when significant body weight loss was observed. Sterile drinking water in a bottle was available to all animal ad libitum during the quarantine and study periods.

The MDA-MB231 tumor cell lines were be maintained *in vitro* as monolayer in L-15 medium supplemented with 10% heat inactivated fetal bovine serum at 37 °C in an atmosphere of 100% air. The tumor cells were routinely sub-cultured before confluence by trypsin-EDTA treatment, not to exceed 4-5 passages. The cells growing in an exponential growth phase were harvested and counted for tumor inoculation. Each mouse was inoculated subcutaneously on the right flank with MDA-MB-231 tumor cells (1 × 10^7^) in 0.1 mL medium and matrigel mixture (1:1 ratio) for tumor development. All animals were monitored for tumor growth. In addition, all animals were monitored for irregular behavior such as changes in mobility, food and water consumption (by cage side checking only), body weight (BW), eye/hair matting or any other abnormal condition. Any mortality and/or abnormal clinical signs were recorded. The measurement of tumor size was conducted with a caliper and recorded twice a week. Groups and treatments were started when the mean tumor volume reached 168 mm^3^. Based on the tumor volume and body weight, mice were randomly assigned to respective groups such that the average starting tumor size and body weight was the same for each treatment group. Mice were provided with vehicle (5% DMSO/95% (20% HP-β-CD in water)) or IDB-003 dissolved in vehicle by oral gavage. For the group dosed with 3 mg/kg IDB-003, the mice received treatments twice daily for 28 days. The mice dosed at 10 mg/kg were given a 2-day holiday after 11 days of treatment (11 days on, 2 days off) for the 28-day period. To analyze TGI, the following formula was used: TGI = 100*(1-(ΔT/ΔC)) where ΔT/ΔC = (TV treated final-TV treated initial)/ (TV Vehicle final-TV Vehicle initial).

## Methods for chemical synthesis of interdictors Scheme SI-1: Synthesis of IDB-001 and IDB-002

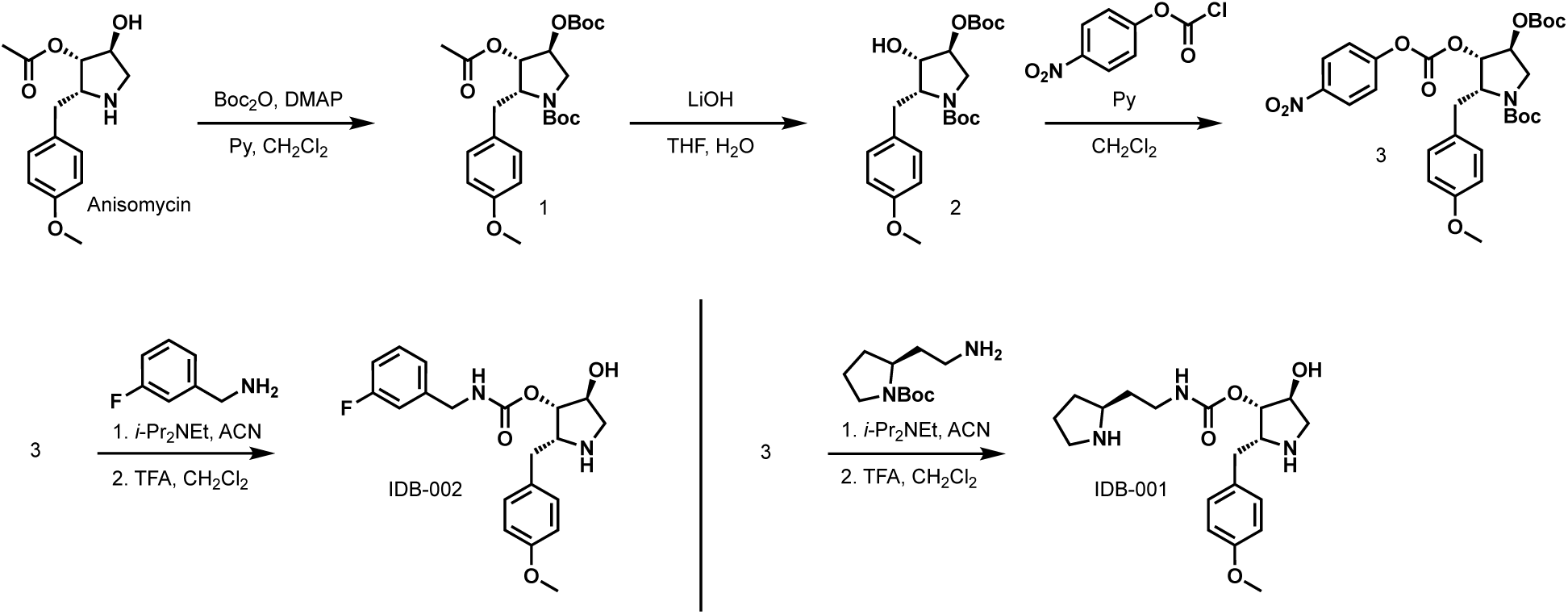

### *Tert*-butyl (2*R*,3*S*,4*S*)-3-(acetyloxy)-4-[(*tert*-butoxycarbonyl)oxy]-2-[(4-methoxyphenyl)methyl]pyrrolidine-1-carboxylate (1)

Boc anhydride (41 g, 188 mmol, 10 equiv) and DMAP (3.45 g, 28.3 mmol, 1.5 eqiv) were added to a solution of anisomycin (5.0 g, 19 mmol, 1.0 equiv) in pyridine (100 mL) at room temperature. After 16 h at 25 °C the reaction mixture was concentrated in vacuo and the residue was purified by reversed-phase flash chromatography (0.05% NH_4_HCO_3_ in H_2_O/ACN) to afford the title compound as a light yellow oil (8.0 g, 91%). MS: m/z: calc’d for C_24_H_35_NO_8_ [M+H-56-100]^+^ = 310, found 310.

### *Tert*-butyl (2*R*,3*S*,4*S*)-4-[(*tert*-butoxycarbonyl)oxy]-3-hydroxy-2-[(4-methoxyphenyl)methyl]pyrrolidine-1-carboxylate (2)

Lithium hydroxide (1.2 g, 51 mmol, 3.0 equiv) and water (10 mL) was added to a solution of **1** (8.0 g, 17 mmol, 1.0 equiv) in THF (100 mL) at room temperature. After 16 h the resulting mixture was extracted with EtOAc (3 x 30 mL) and the combined organic layers were washed with water (3 x 10 mL) and dried over anhydrous Na_2_SO_4_. Concentration under reduced pressure afforded the title compound (7.2 g) as a light yellow solid, which was used directly in the next step without further purification. MS: m/z: calc’d for C_22_H_33_NO_7_ [M+H-56-56]^+^ = 312, found 312.

### *Tert*-butyl (2*R*,3*S*,4*S*)-4-[(*tert*-butoxycarbonyl)oxy]-2-[(4-methoxyphenyl)methyl]-3-[(4-nitrophenoxycarbonyl)oxy]pyrrolidine-1-carboxylate (3)

Pyridine (1.8 g, 22 mmol, 2.0 equiv) was added to a room temperature solution of **2** (4.7 g, 11 mmol, 1.0 equiv) and 4-nitrophenylcarbonochloridate (2.4 g, 17 mmol, 1.5 equiv) in CH_2_Cl_2_ (50 mL) and the mixture was stirred for 2 h. The mixture was concentrated and the resulting residue was purified (SiO_2_, 20% EtOAc in petroleum ether) to afford the title compound as a light yellow oil (5.4 g, 83%). MS: m/z: calc’d for C_29_H_36_N_2_O_11_ [M+H-100]^+^ = 489, found 489.

### (2*R*,3*S*,4*S*)-4-hydroxy-2-(4-methoxybenzyl)pyrrolidin-3-yl (2-((*S*)-pyrrolidin-2-yl)ethyl)carbamate (IDB-001)

A mixture of **3** (67 mg, 0.12 mmol, 1.0 equiv) and *tert*-butyl (2S)-2-(2-aminoethyl)pyrrolidine-1-carboxylate (26 mg, 0.12 mmol, 1.0 equiv) was dissolved in ACN (5 mL) before *i*-Pr_2_NEt (63 µL, 0.36 mmol, 3.0 equiv) was added. The reaction mixture was concentrated in vacuo after 16 h and the residue was used without further purification. The residue was dissolved in 1 mL of 1:1 CH_2_Cl_2_/TFA, the mixture was stirred for 2 h, and concentrated in vacuo. Prep-HPLC purification afforded **IDB-001** (20 mg, 46%, 2 steps).

**^1^HNMR** (400 MHz, Methanol-*d*_4_) *δ* 7.26 – 7.18 (m, 2H), 6.95 – 6.87 (m, 2H), 4.94 (d, *J* = 4.0 Hz, 1H), 4.36 (d, *J* = 4.2 Hz, 1H), 4.18 – 4.14 (m, 1H), 3.78 (s, 3H), 3.61 – 3.47 (m, 2H), 3.34 (d, *J* = 7.7 Hz, 1H), 3.29 – 3.25 (m, *J* = 6.9, 3.0 Hz, 3H), 3.20 – 3.18 (m, 1H), 3.09 (dd, *J* = 14.2, 7.1 Hz, 1H), 2.97 (dd, *J* = 14.1, 8.5 Hz, 1H), 2.32 – 2.24 (m, 1H), 2.17 – 1.84 (m, 4H), 1.74 – 1.64 (m, 1H); **MS** m/z calc’d for C_19_H_29_N_3_O_4_ [M+H]^+^ = 364, found 364.

### (2*R*,3*S*,4S)-4-hydroxy-2-[(4-methoxyphenyl)methyl]pyrrolidin-3-yl *N*-[(3-fluorophenyl)methyl]carbamate (IDB-002)

The title compound was prepared using the same procedure described for IDB-001 with 1-(3-fluorophenyl)methanamine (17 mg, 39% yield, 2 steps).

**^1^H NMR** (400 MHz, Methanol-*d*_4_) *δ* 7.38 – 7.36 (m, 1H), 7.24 – 6.99 (m, 5H), 6.92 – 6.84 (m, 2H), 4.9 (s, 1H), 4.44 – 4.35 (m, 3H), 4.13 – 4.11 (m, 1H), 3.79 (s, 3H), 3.56 (dd, *J* = 12.6, 4.4 Hz, 1H), 3.20 (d, *J* = 12.6 Hz, 1H), 3.09 (dd, *J* = 14.0, 7.7 Hz, 1H), 2.96 (dd, *J* = 14.0, 8.1 Hz, 1H); **MS**: m/z calc’d for C_20_H_23_FN_2_O_4_ [M+H]^+^ = 375, found 375.

## Scheme SI-2: Synthesis of IDB-003

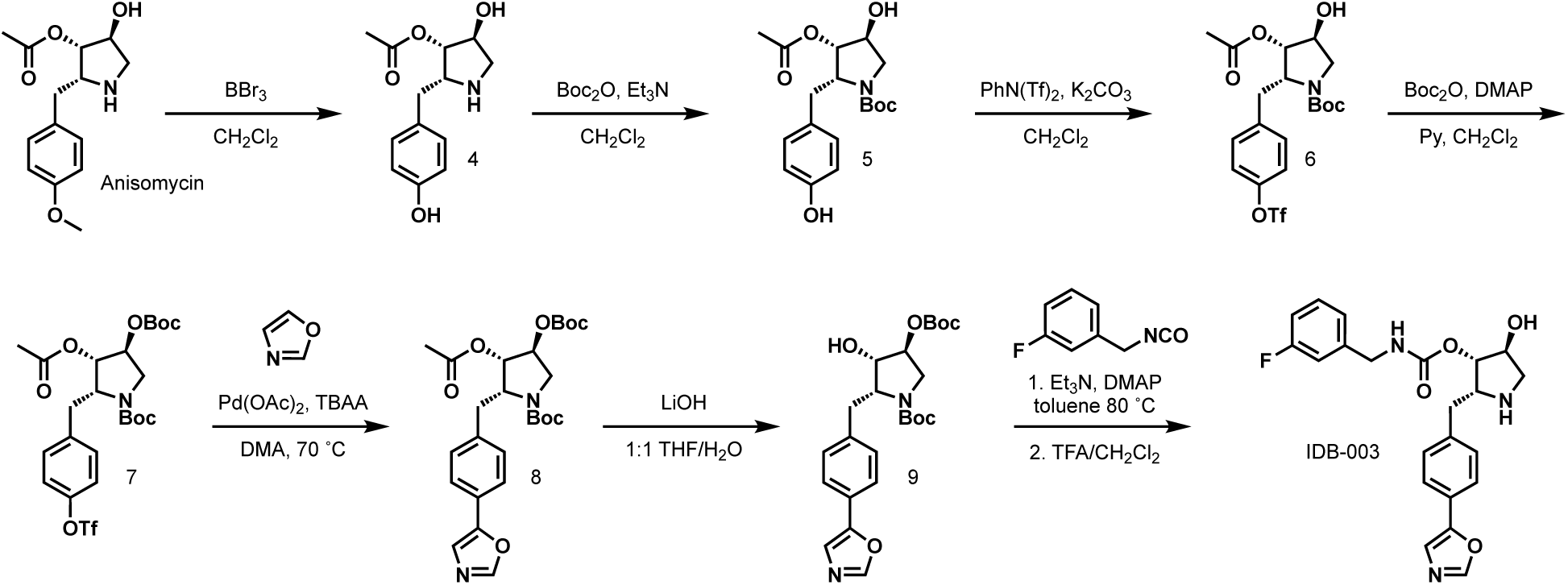

### (2*R*,3*S*,4*S*)-4-hydroxy-2-[(4-hydroxyphenyl)methyl]pyrrolidin-3-yl acetate (4)

A solution of anisomycin (1.5 g, 5.7 mmol, 1.0 equiv) in CH_2_Cl_2_ (8 mL) was cooled to –78 °C before addition of BBr_3_ (16.8 mL, 3 equiv, 1.0M in CH_2_Cl_2_). The reaction mixture was stirred at –78 °C for 2 h, warmed to 25 °C for 1 h, and NaHCO_3_ (sat. aq.) was added. The organic phase was removed and concentration afforded **4** which was used without further purification (1.8 g). MS: m/z: calc’d for C_13_H_17_NO_4_ [M+H]^+^ = 252, found 252.

### *Tert*-butyl (2*R*,3*S*,4*S*)-3-(acetyloxy)-4-hydroxy-2-[(4-hydroxyphenyl)methyl]pyrrolidine-1-carboxylate (5)

A solution of **4** (1.8 g, 7.2 mmol, 1.0 equiv) and Et_3_N (2.5 g, 25 mmol, 3.5 equiv) in CH_2_Cl_2_ (30 mL) was cooled to 0 °C before Boc_2_O (1.9 g, 8.6 mmol, 1.2 equiv) was added and the mixture was warmed to rt for 3 h. The mixture was filtered, the filtrate was concentrated, and reverse-phase column chromatography afforded **5** as a white solid (1.7 g, 68%). MS: m/z: calc’d for C_18_H_25_NO_6_ [M-H]^-^ = 350, found 350.

### *Tert*-butyl (2*R*,3*S*,4*S*)-3-(acetyloxy)-4-hydroxy-2-{[4-(trifluoromethanesulfonyloxy)phenyl]methyl}pyrrolidine-1-carboxylate (6)

A solution of **5** (1.7 g, 4.8 mmol, 1.0 equiv) and K_2_CO_3_ (2.0 g, 14 mmol, 3.0 equiv) in DMF (16 mL) was cooled on an ice bath before the addition of 1,1,1-trifluoro-*N*-phenyl-*N*-((trifluoromethyl)sulfonyl)methanesulfonamide (2.2 g, 6.3 mmol, 1.3 equiv) and the ice bath was removed. After 2 h the reaction mixture was filtered and the filtrate was purified by reverse-phase chromatography to afford **6** (1.5 g, 64%) as a white solid. MS: m/z: calc’d for C_19_H_24_F_3_NO_8_S [M+NH_4_]^+^ = 501, found 501.

### *Tert*-butyl (2*R*,3*S*,4*S*)-3-(acetyloxy)-4-[(tert-butoxycarbonyl)oxy]-2-{[4-(trifluoromethanesulfonyloxy)phenyl]methyl}pyrrolidine-1-carboxylate (7)

A mixture of DMAP (500 mg, 4.1 mmol, 2.0 equiv) and **6** (1.0 g, 2.1 mmol, 1.0 equiv) was dissolved in pyridine (10 mL) and Boc_2_O (4.5 g, 21 mmol, 10 equiv). The resulting mixture was stirred at rt for 24 h and concentrated under reduced pressure. Reverse-phase chromatography afforded **7** (1.0 g, 83%) as a yellow solid. MS: m/z: calc’d for C_24_H_32_F_3_NO_10_S [M+22]^+^ = 606; found 606.

### *Tert*-butyl (2*S*,3*S*,4*S*)-3-(acetyloxy)-4-[(*tert*-butoxycarbonyl)oxy]-2-{[4-(1,3-oxazol-5-yl)phenyl]methyl}pyrrolidine-1-carboxylate (8)

A mixture of **7** (1.0 g, 1.7 mmol, 1.0 equiv), oxazole (180 mg, 2.5 mmol, 1.5 eq.), DMA (10 mL), tetrabutylammonium acetate (770 mg, 2.6 mmol, 1.5 equiv), and Pd(OAc)_2_ (38 mg, 0.17 mmol, 0.10 eq.) was heated to 70 °C for 24 h, cooled, and filtered. The filter cake was washed with THF (2 x 2 mL) and the filtrate was concentrated under reduced pressure. Reverse-phase chromatography afforded **8** as a light yellow solid (660 mg, 77%) as a light yellow solid. MS: m/z: calc’d for C_26_H_34_N_2_O_8_ [M+H]^+^ = 503, found 503.

### *Tert*-butyl (2*S*,3*S*,4*S*)-4-[(*tert*-butoxycarbonyl)oxy]-3-hydroxy-2-{[4-(1,3-oxazol-5-yl)phenyl]methyl}pyrrolidine-1-carboxylate (9)

Lithium hydroxide (160 mg, 6.6 mmol, 5.0 equiv) was added to a solution of **8** (660 mg, 1.3 mmol, 1.0 equiv) in THF (10 mL) and water (1 mL). After 6 h water (10 mL) was added and the pH of the mixture was adjusted to pH = 6 with 1M HCl (aq.). The mixture was extracted with EtOAc (3 x 30 mL) and the combined organic layers were dried (Na_2_SO_4_) and concentrated under reduced pressure. Reverse-phase column chromatography afforded **9** as a light yellow oil (500 mg, 83%). MS: m/z: calc’d for C_24_H_32_H_2_O_7_ [M+H]^+^ = 461, found 461.

### (2*R*,3*S*,4*S*)-4-hydroxy-2-{[4-(1,3-oxazol-5-yl)phenyl]methyl}pyrrolidin-3-yl *N*-[(3-fluorophenyl)methyl]carbamate (IDB-003)

A solution of **9** (51 mg, 0.11 mmol, 1.0 equiv), 1-fluoro-3-(isocyanatomethyl)benzene (25 mg, 0.16 mmol, 1.5 equiv), Et_3_N (46 µL, 0.33 mmol, 3.0 equiv), and DMAP (13.4 mg, 0.11 mmol, 1.0 equiv) in toluene (3 mL) was heated to 80 °C overnight. The mixture was concentrated in vacuo and purified by reversed-phase chromatography. The residue was dissolved in 1 mL of 1:1 CH_2_Cl_2_/TFA, the mixture was stirred for 2 h, and concentrated in vacuo. Prep-HPLC purification afforded **IDB-003** (5.6 mg, 12%, 2 steps).

**^1^H NMR** (400 MHz, Methanol-*d*_4_) *δ* 8.28 (s, 1H), 7.70 (d, *J* = 8.1 Hz, 2H), 7.53 (s, 1H), 7.48 – 7.31 (m, 3H), 7.27 – 7.14 (m, 1H), 7.13 –7.08 (m, 1H), 7.07 – 6.97 (m, 1H), 4.96 (d, *J* = 3.5 Hz, 1H), 4.64 – 4.10 (m, 4H), 3.78 – 3.54 (m, 1H), 3.26 – 3.23 (m, 1H), 3.23 – 3.17 (m, 1H), 3.11 – 3.09 (m, 1H). **MS**: m/z calc’d for C_22_H_22_FN_3_O_4_ [M+H]^+^ = 412, found 412.

## ACKNOWLEDGEMENTS

The authors thank all members of the Interdict Bio team, as well as Jaime Fraser and Danica Fujimori for their helpful discussions, and for feedback during manuscript preparation. We thank David Bulkley and Glenn Gilbert at the UCSF EM core facility for support during sample screening and Alexandrea Shiflett at the University of Michigan LSI cryo-EM facility for support during data collection. We thank Ben Shambaugh (Mindcentric) for IT support. Finally, we’d like to thank the teams at MBC BioLabs and Lilly Gateway Labs where much of this work was completed for their valuable operational support. We also gratefully acknowledge the invaluable input of past and current scientific advisors Alan Ashworth, Erica Bradshaw, Brian Cox, Dean Felsher, Gondi Kumar, Andrea Local, Donald McDonnell, Pamela Munster, and Tom Rovis.

## AUTHOR CONTRIBUTIONS

P.D.D., P.V.S., S.S.B., L.G.H., and A.P.S. designed the study.

P.D.D. and L.F. performed ribosome profiling experiments and analysis.

P.V.S. performed cryo-EM sample preparation, data analysis, and structure determination. M.H., C.S-S., and N.M.B. performed *in vitro* biochemical characterization.

G.W.W., H.K.A., N.M.B., and J.S.S. performed cell-based characterization. All authors analyzed data and participated in discussions on content.

P.D.D., P.V.S., M.H, S.S.B., L.G.H, and A.P.S. wrote the manuscript with input from all authors.

## EXTENDED DATA FIGURES & FIGURE LEGENDS

**Extended Data Figure 1.**
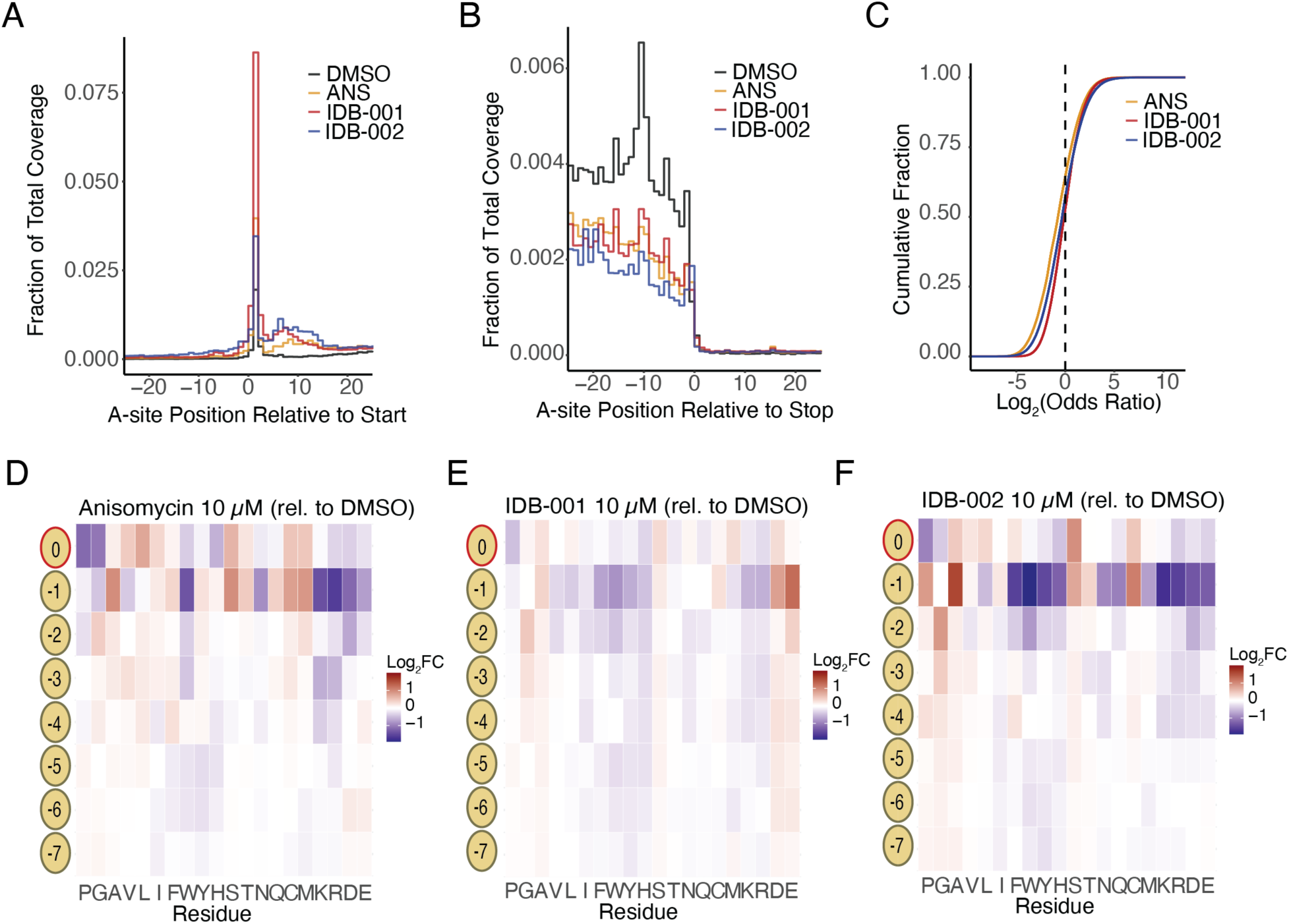
Interdictors lead to unique context-dependent translation inhibition in human cells. A. Metagene analysis of region near start codons for IDB and DMSO-treated samples (codons). B. Metagene analysis of region near stop codons for IDB and DMSO-treated samples (codons). C. Cumulative distribution function of odd-ratios (log2) for codon positions across IDB-001, IDB-002 and ANS treatments relative to DMSO. D-F. Heatmap demonstrating changes in ribosome occupancy for each position in the nascent chain and amino acid residue identity between anisomycin and DMSO (D), IDB-001 and DMSO (E), or IDB-002 and DMSO (F). Red color indicates a log_2_ fold change of +1, and blue –1.

**Extended Data Figure 2.**
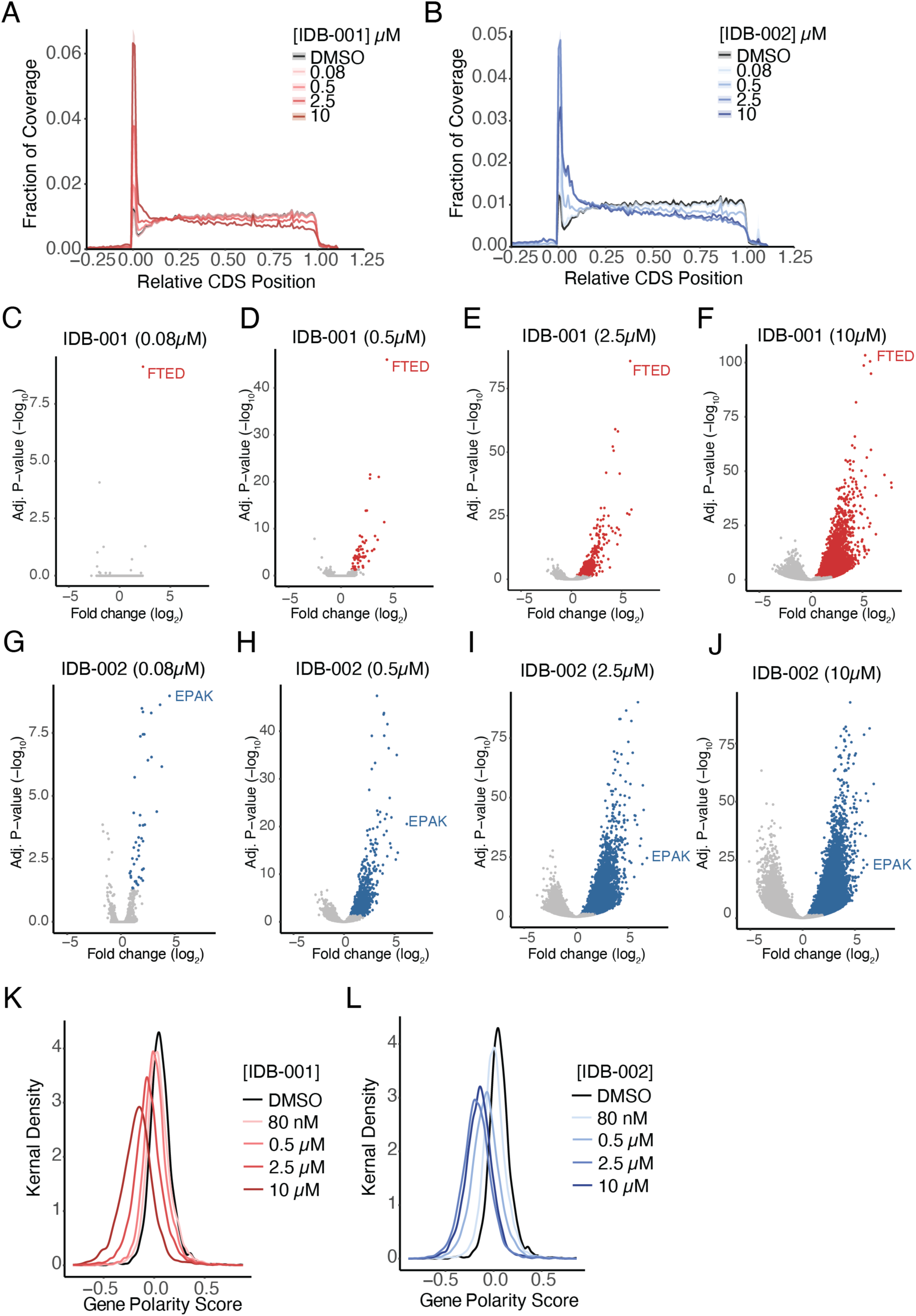
Global and specificity profiles of interdictors are concentration dependent. A-B. Metagene analysis of open-reading frames for IDB-001 (A) and IDB-002 (B) and DMSO-treated samples at various concentrations. C-J. Volcano plots highlighting significantly enriched 4-mer sequences (nascent chain positions – 3 to 0) for IDB-001 (C-F) and IDB-002 (G-J) at various concentrations. K-L. Kernel density plot of polarity scores across transcripts with sufficient coverage (> 0.1 read/codon) for IDB-001 (K) and IDB-002 (L).

**Extended Data Figure 3.**
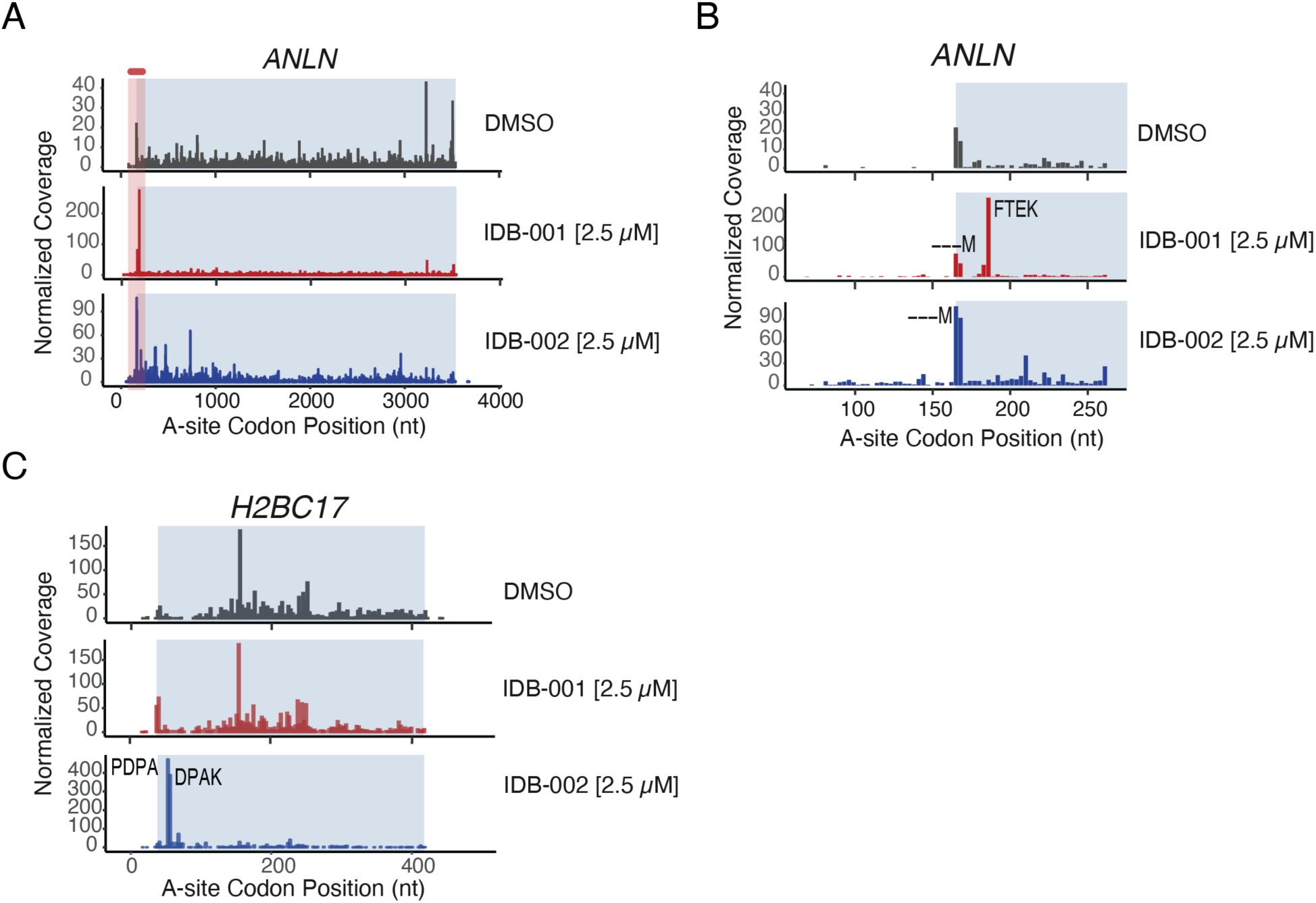
Representative examples of ribosome stalling induced by interdictors on endogenous genes. A-B. Ribosome footprints (adjusted to A site) over the *ANLN* locus (A) and a zoomed into relevant region within *ANLN* locus (B) for cells treated with either DMSO or 2.5 µM IDB-001 or IDB-002. Read counts were scaled based on library size. Blue color signifies the main open reading frame. Novel peaks are labeled with corresponding nascent peptide sequence (positions –3 to 0). C. Same as (A) over the *H2BC17* locus.

**Extended Data Figure 4.**
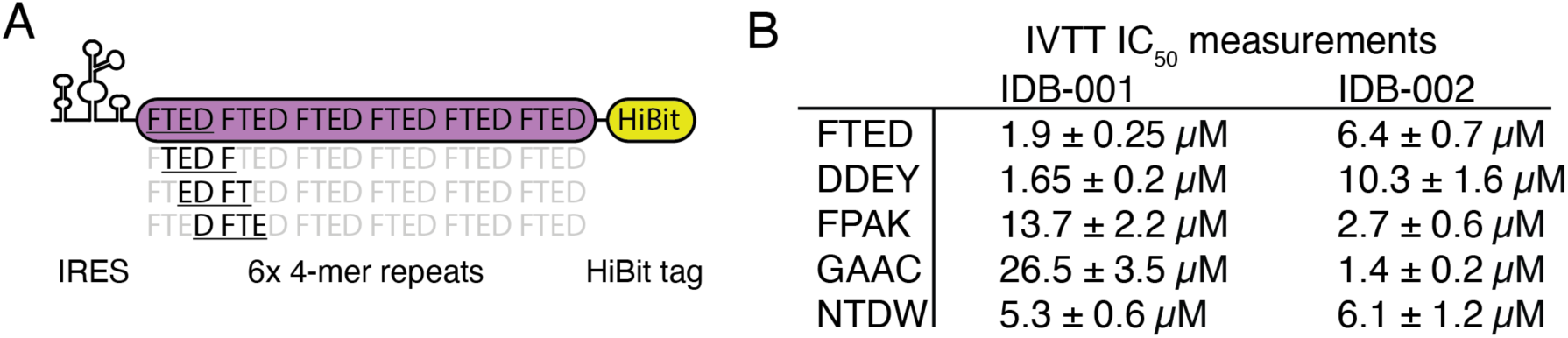
*In vitro* kinetic analysis of interdictor translation inhibition. A. Schematic representation of the HiBit containing reporter constructs illustrating the different 4-mers encountered by the interdictors as the ribosome proceeds down the mRNA. B. Table of the IC_50_ values determined for each interdictor-reporter combination shown in Figure 2E. Errors are SEM (n=4 experiments).

**Extended Data Figure 5.**
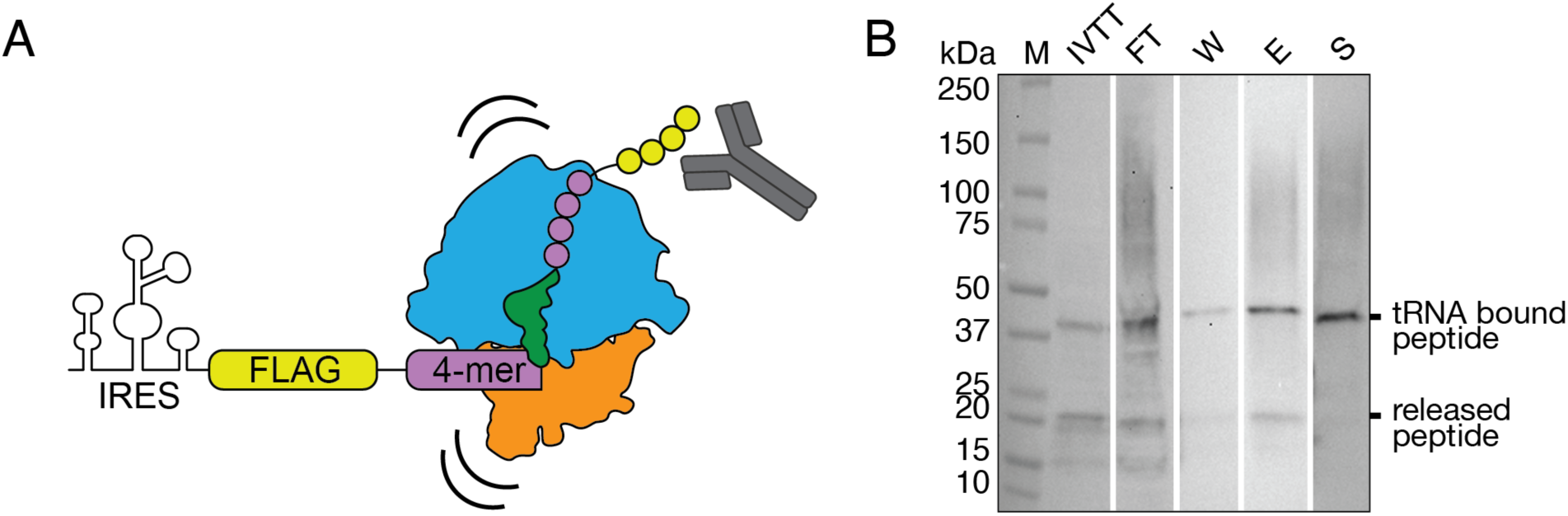
Preparation and purification of 4-mer containing human RNCs. A. Schematic showing construct design and purification approach. Ribosomes stalled on the last codon of a non-stop mRNA can be purified by anti-FLAG pulldown. B. Anti-FLAG Western blot showing steps from RNC purification. Relevant lanes (from the same blot) are shown: IVTT – soluble fraction of the *in vitro* transcription translation reaction after 20 min. FT – unbound fraction after incubation with FLAG beads. W – wash fraction. E – elution fraction. S – sample for cryo-EM after concentration, buffer exchange and size selection. FLAG band at ∼40 kDa corresponds to a FLAG-peptidyl-tRNA species. When cleaved or released from tRNA, the peptide has a size of ∼20 kDa.

**Extended Figure 6.**
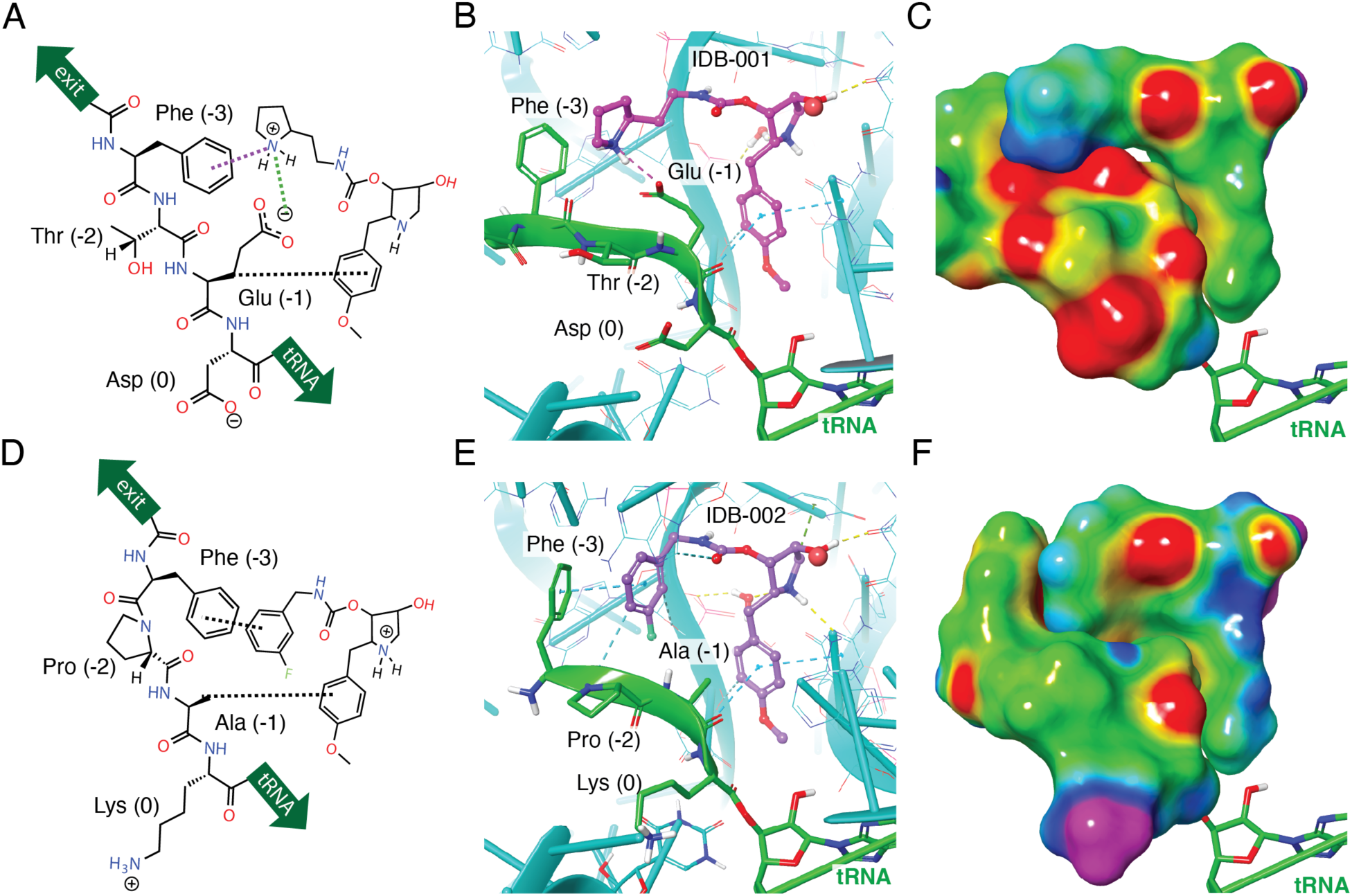
IDB-001 and IDB-002 make specific interactions with nascent polypeptide. A. Schematic representation of the relative orientation and interactions between IDB-001 and the nascent polypeptide chain as observed by cryo-EM. B. Model for structure of IDB-001 (pink) bound to FTED peptide highlighting key interactions with the sidechains of the FTED peptide. rRNA (blue) and nascent peptide (green) are shown within 6 Å of IDB-001.). C. Electrostatic potential surface representation of (B) to highlight the shape and charge complementarity of IDB-001 with FTED peptide. Coloring is as follows: red/orange indicates electron-rich, negative charge; purple/blue indicates electron-deficient, positive charge; green indicates neutral). Images, electrostatic potential maps and surfaces generated in Maestro (Schrödinger Release 2024-2: Maestro, Schrödinger, LLC, New York, NY, 2024.) D. Same as in (A) for IDB-002. E. Same as in (B) for IDB-002 with the FPAK peptide. F. Same as in (C) for IDB-002 with the FPAK peptide.

**Extended Data Figure 7.**
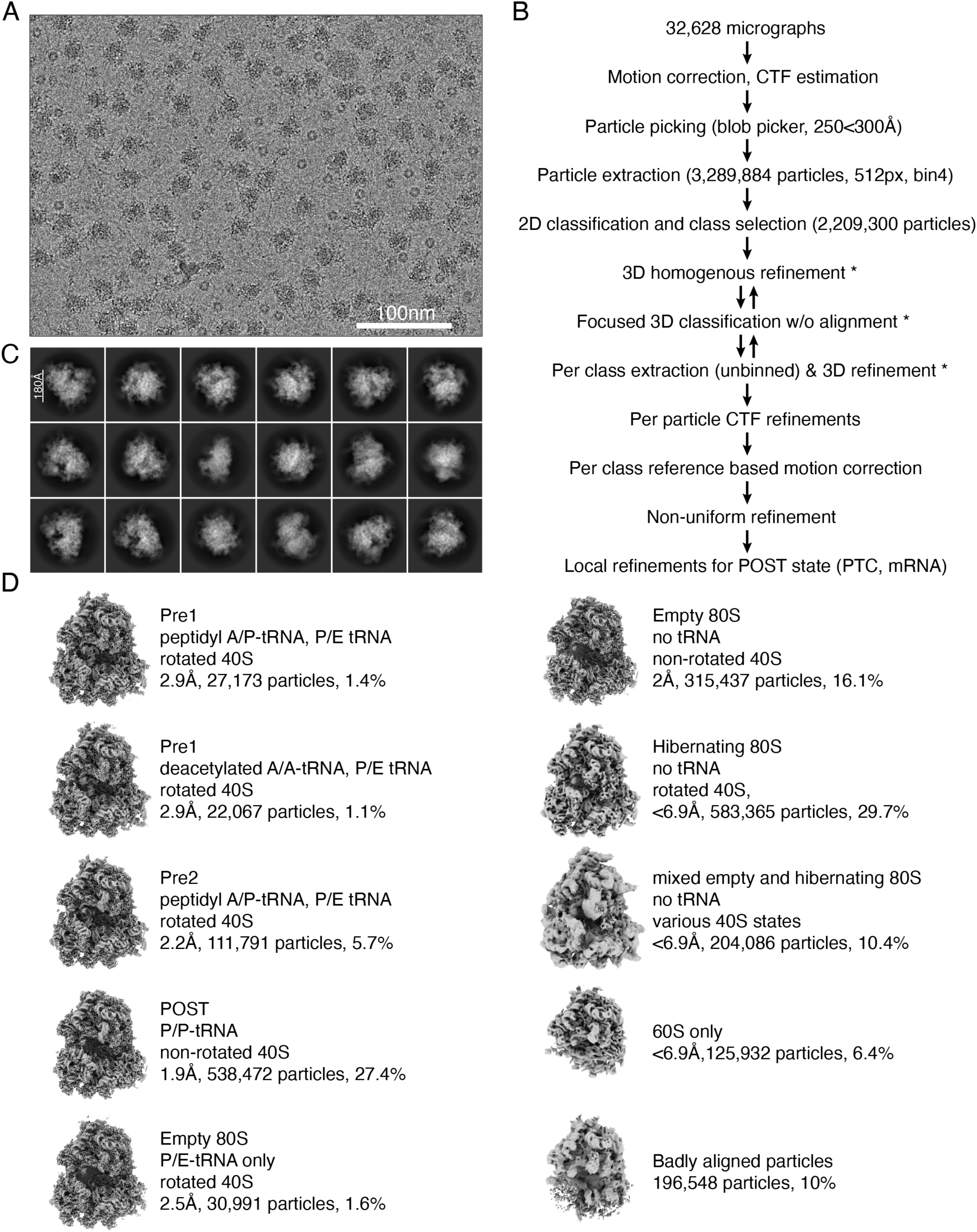
Representative cryo-EM processing workflow for the stalled MYC:IDB-001 RNC. A. Representative cryo-EM micrograph. 32,628 similar micrographs were collected. B. Processing workflow. At steps marked with an asterisk, major classes shown in D were identified. C. Representative 2D class averages showing different views of the ribosome particles D. Identified classes form the MYC:IDB-001 RNC dataset. Classes are distinguished by functional state, subunit composition and tRNA presence. Map resolution (FSC = 0.143), particle numbers and percentage of total number of particles are indicated next to each class.

**Extended Data Figure 8.**
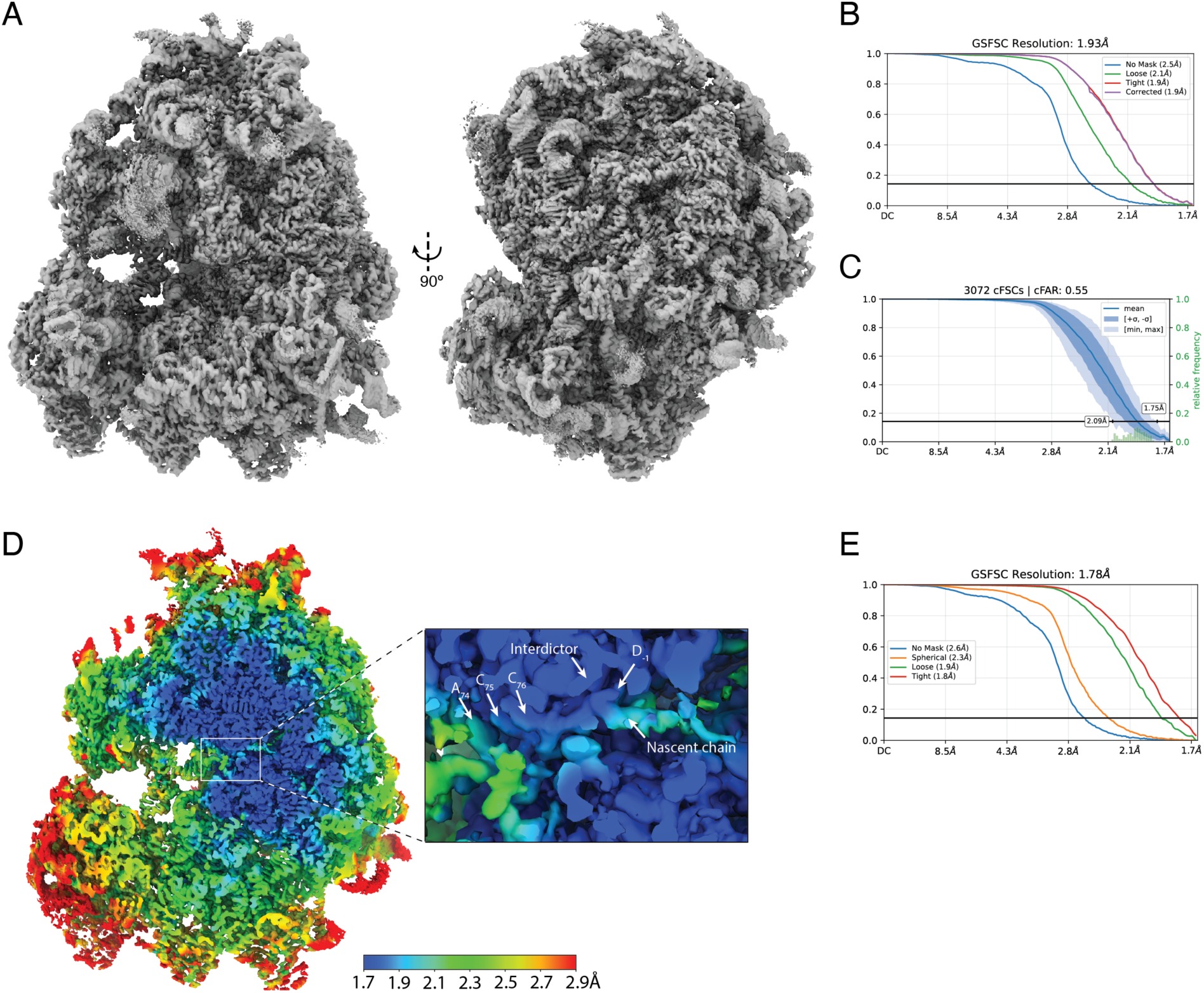
Cryo-EM map validation. A. Consensus refinement map for the stalled MYC:IDB-001 RNC complex with a peptidyl P/P tRNA. B. FSC curve for (A). C. Conical FSC distribution for A, showing homogenous spherical distribution of resolution and the absence of major preferred orientation, as implemented in CryoSPARC v.4.6.2. D. Local refinement map that focuses on the PTC and that is colored by local resolution (in angstrom). Inset shows the PTC with the interdictor, the nascent chain, and the tRNA CCA indicated by arrows. E. FSC curve for (D).

**Extended Data Figure 9.**
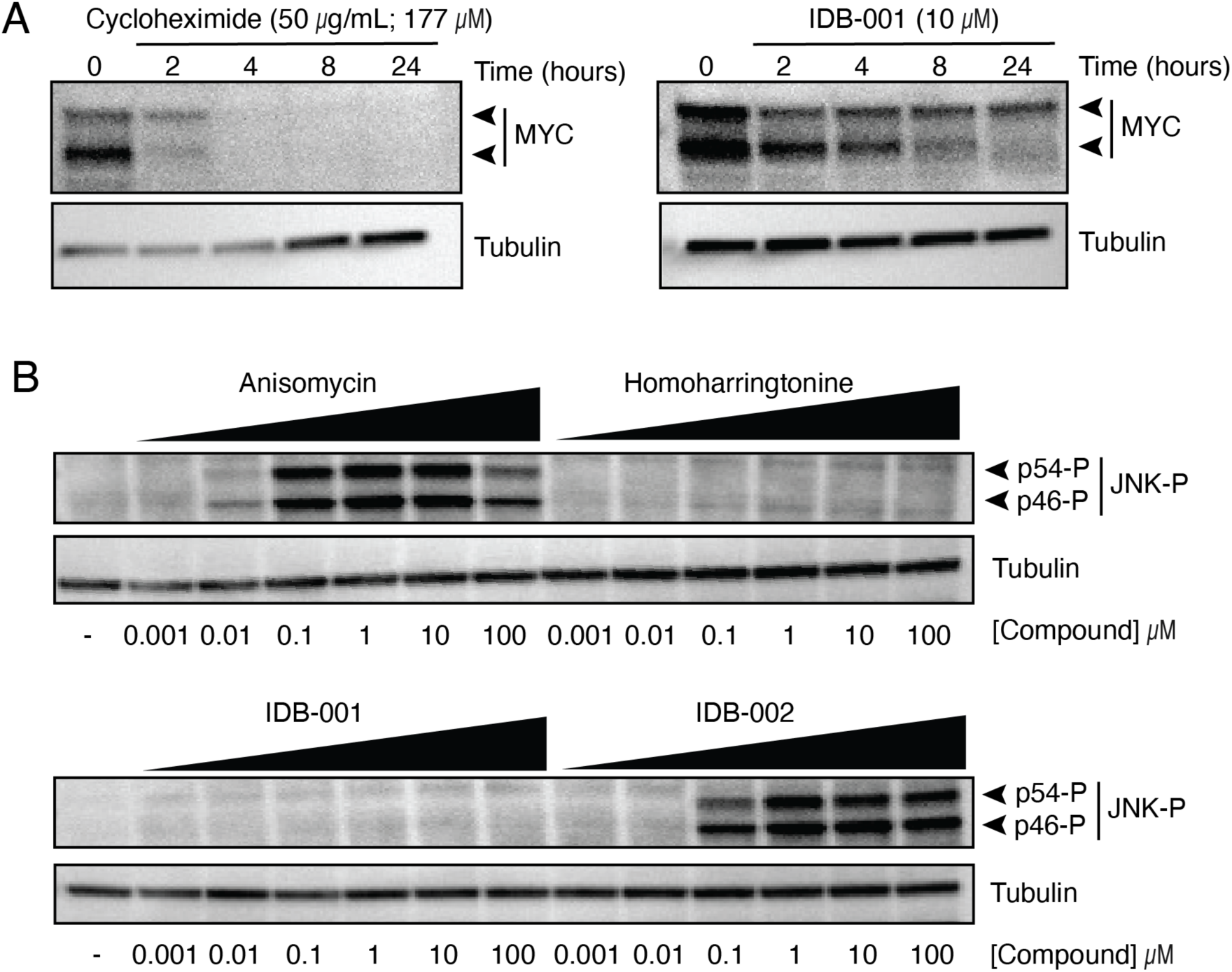
Interdictors lead to MYC depletion and differentially regulate the RSR. A. Western blot analysis of MYC protein in HCC-1143 triple-negative breast cancer cells treated with either cycloheximide (50 µg/mL) or IDB-001 (10 µM). B. Western blot analysis for JNK phosphorylation (JNK-P), a marker of ribotoxic stress pathway with varying interdictor or translation inhibitor treatment concentrations in HEK293T cells.

**Extended Data Figure 10.**
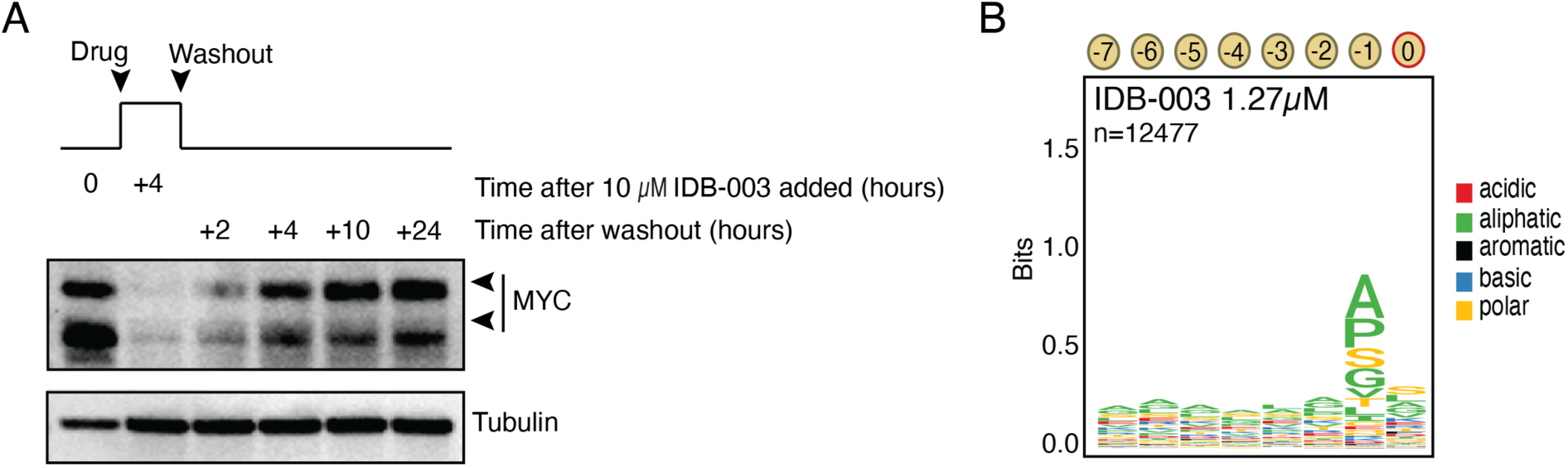
IDB-003 shows similar context-dependent activity as IDB-002 and leads to MYC depletion in HCC-1143 cells. A. Western blot analysis of MYC protein in HCC-1143 cells treated with IDB-003 (10 µM) for 4 hours, then the media was replaced to wash out drug, and MYC resynthesis observed over an additional 24-hour period. B. Sequence logo for the nascent chain sequence from positions –7 to 0 for significantly enriched pause sites codons, as in Figure 1E-G, but for IDB-003 (1.27 µM).

